# Inflammatory injury drives CIN and STING loss in polyploid pancreatic acinar cells

**DOI:** 10.64898/2026.02.10.704772

**Authors:** Jan Brunken, Enrico Frigoli, Azmal Syed Ali, Veronika Elisabeth Lummer, Axel Carbone, Andrea Ruiz Sarvari, Sefa Berkay Çayır, Sanzhar Aitbay, Daria Kocherhina, Sonja Anslinger, Katharina Bauer, Ulrike Engel, Jan-Philipp Mallm, Eli Pikarsky, Simon Anders, Jeroen Krijgsveld, Ana Martin-Villalba

## Abstract

Epidemiological studies link pancreatic inflammation to increased cancer risks with elevated levels of chromosomal instability (CIN), yet the underlying cause remains unclear. The exocrine pancreas and lactating mammary gland harbor polyploid cells to boost secretory efficiency. Using organoid and *in vivo* models of pancreatic injury, we show that polyploid acinar cells contribute to regeneration via acinar-to-ductal metaplasia. The metaplasia induced cell size reduction is associated with crowding and spindle misorientation, leading to mitotic errors such as lagging chromosomes, chromatin bridges, and micronuclei. Micronucleated cells are not always eliminated or arrested in the cell cycle but can continue to proliferate and expand. Polyploid cells in post-lactational mammary gland organoids equally exhibited proliferation-related CIN. Single-cell proteomics of acinar cells shows STING1 loss and increased ATR levels in polyploid cells, enabling continued proliferation. These findings uncover physiological links between polyploidy, tissue repair, and CIN, offering new insights into cancer risks after tissue remodeling.

## Introduction

Epidemiological and experimental evidence strongly implicates inflammation and regenerative responses as major contributors to cancer development, particularly in tissues subject to chronic injury, repair or remodeling such as glandular organs like the pancreas, liver and mammary glands^1–6^. Interestingly, these organs contain significant amounts of binucleated polyploid cells, which have largely been overlooked in the context of regeneration and tumor initiation^7–9^. These cell types may represent a unique, endogenous source of chromosomal instability, particularly under conditions of cellular stress or tissue repair. Polyploidization, while often viewed as a mechanism to enhance cellular resilience, may increase the risk of mitotic errors, leading to aneuploid progeny and genomic instability as it is well established in various cancer types^10–13^. Despite their presence in malignant tumors, the role of naïve polyploid cells in initiating spontaneous chromosomal instability (CIN) remains poorly defined.

CIN is a hallmark of many cancers and a major driver of tumor evolution, observed in over 80% of solid tumors^14–18^. Defined by an increased rate of chromatin gains or losses due to chromosome mis-segregation, CIN is closely linked to aneuploidy, micronuclei formation, chromothripsis, and widespread structural variants, features that collectively contribute to elevated mutational burden^19–21^. Notably, certain cancer types such as pancreatic ductal adenocarcinoma (PDAC), mammary gland tumors, and hepatocellular carcinoma (HCC) exhibit particularly high levels of CIN, often arising at early stages of tumorigenesis including pre-malignant and pre-invasive lesions^22–24^. Importantly, pancreatic intraepithelial neoplasia (PanIN) lesions harbor chromothriptic events in approximately 40% of evaluated lesions, underscoring the early emergence of complex genomic rearrangements in PDAC^25^. While CIN is a well-established feature of tumor progression, resistance and relapse, the precise initiating events that lead to its emergence remain poorly understood^26–28^. Most research to date employs genetic manipulation or exogenous mutagenic factors, such as radiation, chemical carcinogens or mitosis-perturbing reagents as triggers of genomic instability^29^. While these approaches have been instrumental in describing CIN generation mechanistically and elucidating its impact in the cell’s physiology, they do not explain how CIN is unleashed under physiological conditions. Thus, one substantial question remains: what intrinsic, cell-autonomous mechanisms might predispose cells to CIN in the frame of inflammation linked to regeneration or tissue remodeling events?

In this study, we use *in vivo* and organoid models of the untransformed regenerating mouse pancreas to investigate the regenerative potential of binucleated pancreatic acinar cells as well as their putative role in CIN accumulation. Our findings reveal that these cells, contrary to their previously understood static nature, actively participate in pancreatic regeneration. During this process, binucleated pancreatic acinar cells acquire micronuclei with extensive DNA damage indicating chromothripsis-like events – a phenomenon involving the catastrophic shattering of chromosomes. Intriguingly, we find that these cells do not inevitably undergo apoptosis or senescence but often survive and even remain proliferative without further treatment or modification. We further show that these cells downregulate STING1 and upregulate ATR as a CIN adaptation response. By linking these cellular events to regenerative processes, our study provides new insights into how polyploid epithelial cells may serve as a source of mutations that drive pancreas and breast carcinogenesis.

## Results

### Binucleated pancreatic acinar cells actively participate in tissue regeneration

We previously reported that more than 40% of pancreatic acinar cells in mice and 15% in humans are binucleated^7^. Furthermore, we concluded that binucleated acinar cells would represent a terminally differentiated cell state, not contributing to proliferation during homeostasis or injury^7^. However, the effect of pancreatic injury on the number of nuclei and ploidy of the acinar population was not fully assessed. Thus, we first asked if a pancreatic injury alters the mononucleated-to-binucleated cell ratio and whether this shift is reversed upon regeneration. To examine this, we employed cerulein-induced pancreatitis (CiP) in mice and fixed pancreas tissue at different days after the last cerulein injection (dpi)^30^. We assessed the presence of binucleated cells over the course of an acute inflammatory response (2 dpi, 4 dpi) to the fully regenerated organ (28 dpi to 91 dpi, Figure 1A). We found that the fraction of binucleated acinar cells drops from 40% to approximately 32% at 2 dpi and 24% at 4 dpi (Figure 1B, C). This fraction rose to 31% at 28 dpi and was almost back to original value of 40% at 91 dpi. Thus, acute inflammation leads to a decreased number of binucleated cells that is restored as regeneration is complete, in line with the proposed role of binucleated cells in liver size control^31^.

**Figure 1.**
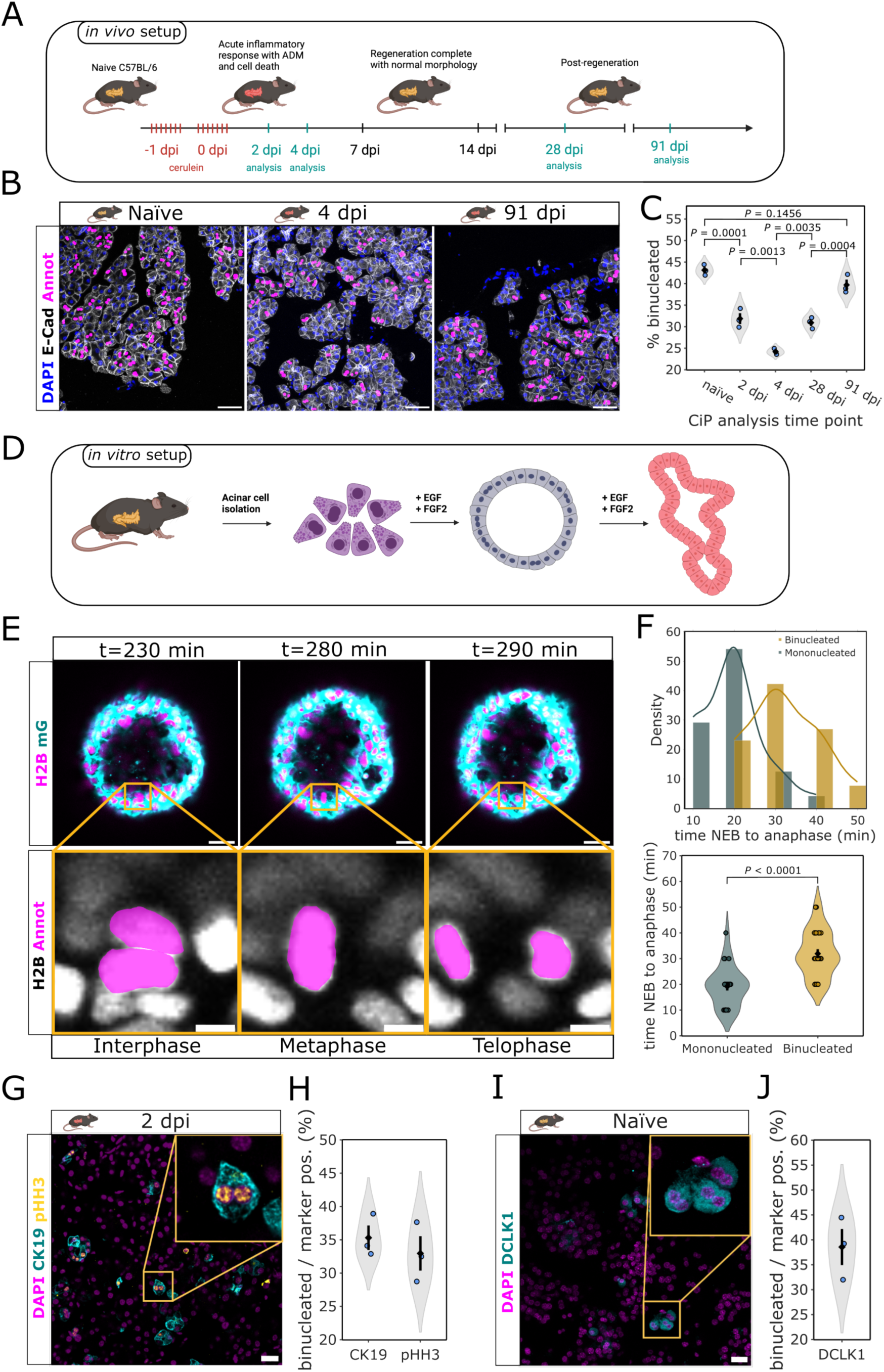
Polyploid pancreatic acinar cells contribute pancreas regeneration. (A) Experimental setup to assess pancreas regeneration and post-regeneration after acute inflammation using cerulein-induced pancreatitis. Scheme created in Biorender. (B) Immunofluorescence (IF) staining of healthy (naïve), regenerating (4 dpi) and post-regenerative (91 dpi) mouse pancreatic tissue. Injury was mediated by cerulein-induced pancreatitis (CiP). DAPI (blue), E-Cadherin (white) and annotations indicating binucleated cells (magenta). Scale bar: 50 μm. (C) Quantification of the fraction of binucleated acinar cells in naïve and injured (2 dpi, 4 dpi and 91 dpi) mouse pancreata. *P* values were calculated by one-way ANOVA followed by Tukey’s post hoc test (n = 3 mice). Violin plots show the distribution of individual biological replicates; black diamonds indicate the mean and black bars indicate s.e.m. (D) Experimental setup to assess the role of polyploid pancreatic acinar cells in regeneration and early tumor formation using mouse acinar-derived organoids (mADOs). Scheme created in Biorender. (E) Live-cell image series showing a binucleated acinar cell mitosis in H2B-mCherry/mG mice in interphase, prophase and telophase. H2B (magenta), plasma membranes/mG (cyan). Scale bar: 20 μm. Yellow squares highlight zoomed regions with H2B (white) and binucleated cell annotation (magenta). Scale bar: 5 μm. (F) Duration measurements from nuclear envelope breakdown to anaphase derived from mitotic cells from live-cell imaging data. Upper graph shows density distribution and kernel density plot of mononucleated (cyan) and binucleated (yellow) cell durations. Lower panel shows violin plots of mononucleated (cyan) and binucleated (yellow) cell durations. *P* value was calculated using a two-sided unpaired Student’s *t* test (n = 50 mitotic events from 3 mice). (G) IF image of injured mouse pancreas (CiP, 2 dpi) to assess the contribution of binucleated acinar cells to ADM (cytokeratin 19-expressing, CK19) and proliferation (phospho-histone H3-positive, pHH3). DAPI (magenta), CK19 (cyan), pHH3 (yellow). Yellow box indicates zoomed region. Scale bar: 20 μm. (H) Quantification of binucleated cell fraction in pHH3- and CK19-expressing acinar cells in cerulein-induced pancreatitis at 2 dpi. *P* value was calculated using a two-sided unpaired Student’s *t* test (n = 3 mice). (I) IF image of naïve mouse pancreatic tissue to assess the contribution of binucleated acinar cells to the pool of facultative progenitors (Doublecortin-like kinase 1-expressing, DCLK1). DAPI (magenta), DCLK1 (cyan). Scale bar: 20 μm. (J) Quantification of binucleated cell fraction in DCLK1-expressing naïve acinar cells. *P* value was calculated using a two-sided unpaired Student’s *t* test (n = 3 mice).

We next examined whether binucleated cells are generated through cell fusion or endomitosis after acute inflammation. To this end, we administered bromodeoxyuridine (BrdU) immediately after the last cerulein injection. At 91 days, no binucleated cells with mixed BrdU^+^/BrdU^−^ nuclei were observed. Instead, every BrdU-labeled binucleated cell had both nuclei labeled, favoring a mechanism involving endomitosis as the main mode of regeneration (Figure S1A). Considering the short window of BrdU bioavailability of several hours^32^ after the last cerulein injection, this furthermore shows that binucleated acinar cells are generated by cells that were cycling as an early response to injury, at a time when the overall amount of binucleated cell is still decreasing.

Next, we investigated the loss of binucleated cells during the early phase of regeneration. Since binucleated acinar cells do not exhibit an increased propensity for apoptosis following pancreatic injury ^7^, the reduction in the binucleated cell proportion cannot be solely attributed to cell death. An alternative explanation may be the division of binucleated cells into mononucleated cells. To follow the fate of binucleated acinar cells in real time, we utilized an *in vitro* 3D culture of primary mouse acinar-derived organoids^7^ (mADOs, Figure 1D). This model recapitulates acinar-to-ductal metaplasia (ADM) and tubular structure formation, processes that acinar cells undergo *in vivo* to support proliferation and pancreatic regeneration^33,34^ (Figure S1B). First, mADOs were fixed at different time points in culture and stained for E-Cadherin to quantify cell nuclei counts and ploidy. We additionally stained for α-Amylase to assess ADM (Figure S1C). As reported previously, the number of α-Amylase expressing cells decreased during mADO culture indicating the onset and progression of ADM^7^ (Figure S1D). In accordance with our *in vivo* pancreatitis data, the fraction of binucleated cells in mADOs decreased from ∼40% to approximately 13% at 13 days (d13) of culture (Figure S1E). Interestingly, the overall number of polyploid cells did not change proportionally but dropped from 42% at d0 to 30% at d13.

Secondly, we tracked the fate of binucleated cells in mADOs in real time by live confocal and light-sheet imaging over several hours. To distinguish between mono- and binucleated cells we took advantage of H2B-mCherry/mG double reporter mice which fluorescently label the cell nuclei as well as plasma membranes in all cells^35^. Intriguingly, we observed that binucleated acinar cells do divide (Figure 1E, Movie S1). During mitosis, both nuclei condense into a single metaphase plate and give rise to two mononucleated cells. While the binucleated mitoses exhibited otherwise normal mitotic phases, the time from nuclear envelope breakdown (NEB) to anaphase and therefore the overall duration of cell division was significantly prolonged (Figure 1F). This observed delay presumably stems from a prolonged spindle assembly checkpoint during metaphase that ensures attachment of all kinetochores to microtubule fibers. To further examine whether the proliferative binucleated cells perform a full cell cycle, including an S phase, or just represent a diploid cell state locked at a late cell cycle stage, we performed short-term BrdU labeling of mADO cultures starting directly after cell plating. We labelled cells in fresh wells every 2 h and fixed after a pulse time of another 2 hours. We began to see BrdU^+^ cells as early as 24 h after cell plating, including binucleated cells (Figure S1F). In all cases, both nuclei were always BrdU^+^, as previously shown in pancreatitis *in vivo*. These results indicate that binucleated cells represent a true polyploid cell state, capable of proliferating with synchronized onset of replication in both nuclei. In contrast, our previous quantification of prophase cells by phospho-histone H3 (pHH3) staining at 5 days after pancreatitis induction, fail to detect any actively cycling binucleated cell^7^. Thus, this time we quantified proliferating (pHH3^+^) and transdifferentiating (CK19^+^) acinar cells at an earlier time point of 2 dpi. To distinguish mononucleated from binucleated pHH3^+^ cells, we focused our quantification on late G2- and early M-phase nuclei, indicated by speckled pHH3 staining co-localizing with round and not fully condensed nuclear DAPI stain^36^. Notably, we did not only detect binucleated cells that stain positive for CK19 and pHH3, but at 2 dpi approximately 35% of all CK19-expressing and 33% of all pHH3-positive acinar cells were binucleated (Figure 1G, 1H). These findings imply that binucleated acinar cells undergo ADM and proliferate at this early stage of regeneration. It also shows that they have a slightly reduced propensity or slower dynamics to acquire a proliferating ADM state as compared to their mononucleated counterparts. Using long-term lineage multicolor tracing, we previously demonstrated that cerulein-induced injury activates proliferation of a subset of otherwise quiescent acinar cells to replenish lost tissue^7^. Others have reported that this pool of facultative progenitor acinar cells, which undergo ADM upon CiP, can be specifically labeled by doublecortin-like kinase protein 1 (DCLK1)^37,38^. Indeed, we found that approximately 39% of DCLK1 expressing acinar cells are binucleated, supporting a significant contribution of binucleated acinar cells to the subset of facultative progenitor cells.

DCLK1^+^/CK19^+^ binucleated cells could get arrested in G1 in an *in vivo* setting by a “tetraploidy checkpoint” – a potential mechanism to prevent cells from cycling after polyploidization that has been proposed in the early 2000s^39,40^. Although a clearly defined tetraploidy checkpoint has been rejected, recent studies suggest that polyploid hepatocytes can get cell cycle repressed by the PIDDosome/Caspase 2 axis^41,42^. To assess if binucleated acinar cells would fully enter mitosis or rather become arrested after transdifferentiation, we stained mouse pancreatic tissue from cerulein-induced pancreatitis for γ-Tubulin. Polyploidy is usually accompanied by an increased number of centrosomes^43,44^. γ-Tubulin labels nucleation sites for microtubule formation at centrosomes and can thus be used as a proxy for cell ploidy. These extra centrosomes can lead to a multipolar spindle formation with up to 4 individual spindle poles in case of a tetraploid cell with 4 centrosomes^43,45–47^. We screened pancreas tissue after CiP for such metaphase figures that would correspond to polyploid cells. At 2 dpi, we found multipolar metaphases in CiP with up to 8 centrosomes, indicating that tetraploid and even octoploid cells can enter mitosis (Figure S1G). Of note, we found no cell with two metaphase plates, indicating that so called “double mitoses”, in which both nuclei undergo mitosis individually, do not play major role in pancreatic regeneration. This supports that binucleated cells divide via nuclear aggregation as observed in organoids.

### Metaplasia-mediated cell size reduction promotes ploidy reduction in binucleated acinar cells

Multipolar mitoses have been observed in hepatocytes as part of the so-called “ploidy conveyor,” a process that describes the dynamic de- and re-polyploidization to regulate ploidy levels and facilitate adaptive advantages in regenerating liver tissue^48^. However, unlike the regenerating pancreas, hepatocyte proliferation during liver regeneration does not require metaplasia. Hepatocyte-to-cholangiocyte transdifferentiation is not a normal feature of regeneration but rather occurs in response to chronic injury, such as biliary obstruction or liver fibrosis^49^. In contrast, acinar-to-ductal metaplasia (ADM) is essential for acinar cell proliferation and pancreatic regeneration.

Thus, we examined whether the additional constrain imposed by metaplasia in the pancreas influences the tendency to undergo multipolar cell division and ploidy reduction in mononucleated or binucleated polyploid acinar cells. We first screened our H2B-mCherry mADO live-cell imaging data for fully completed multipolar mitoses. We additionally employed live-cell imaging of mADOs of the EGFP-Tuba mouse model, in which fluorescently labeled α-Tubulin enables the visualization of the individual microtubule spindles. We frequently observed mitoses completing with three poles, generating three daughter cells, two of which exhibited reduced DNA content (Figure 2A, Movie S2). We quantified 54 multipolar mitotic events in live cell recordings from 3 EGFP-Tuba mice. Notably, we found that multipolar mitoses predominantly occurred in binucleated cells while mononucleated polyploid cells rarely underwent this process (Figure 2B). Instead, their centrosomes tended to cluster and form two spindle poles.

**Figure 2.**
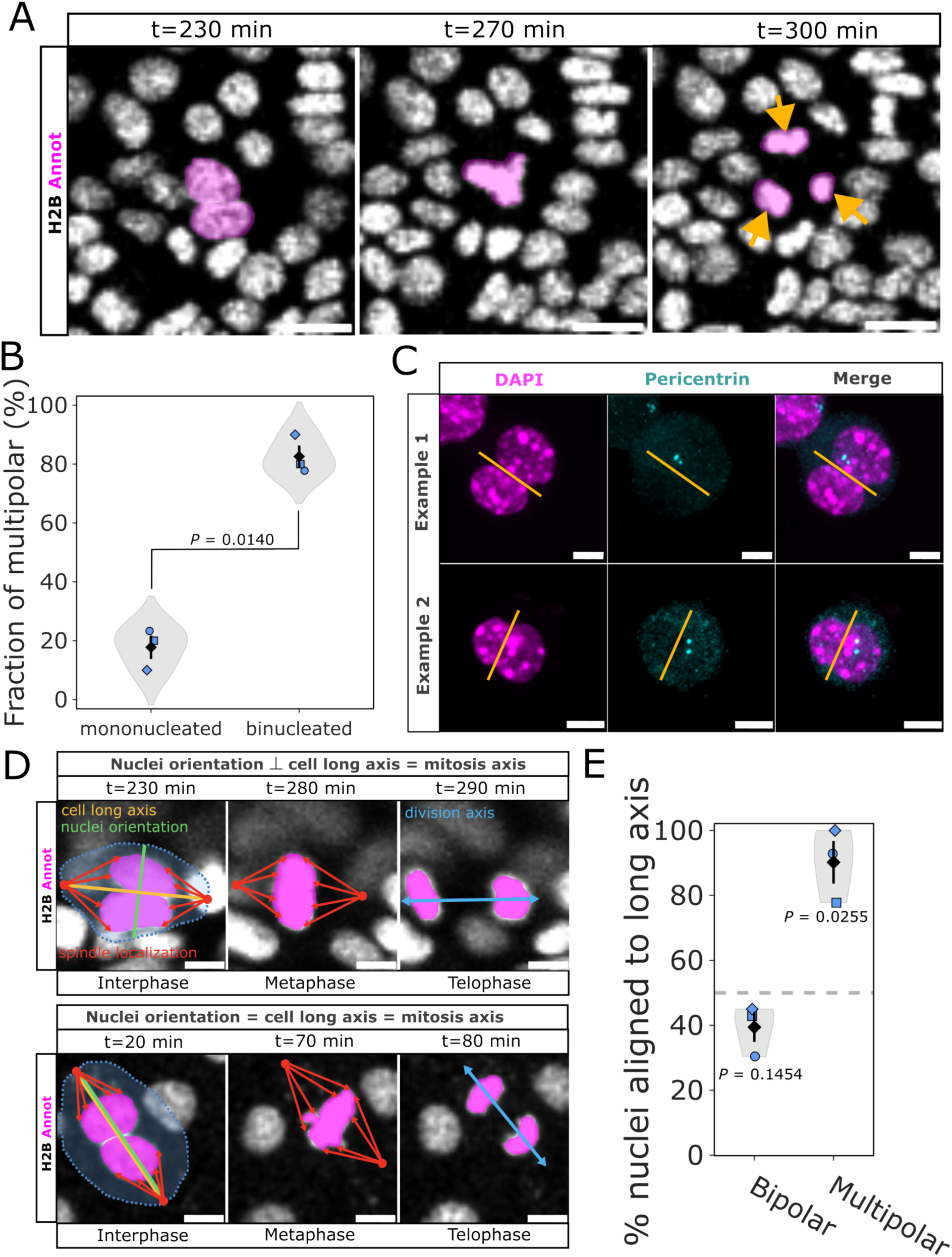
Binucleated acinar cells are predisposed for multipolar spindle orientations. (A) Live-cell image series from H2B-mCherry mADOs showing binucleated cell undergoing multipolar mitosis leading to a ploidy reduction of daughter nuclei. H2B (white), nuclei annotation (magenta). Scale bar 20 μm. (B) Quantification of nuclear numbers of cells undergoing multipolar divisions from live-cell imaging data from EGFP-Tuba mADOs. *P* value was calculated using a two-sided paired Student’s *t* test (n = 3 mice). (C) IF images of two examples (rows) of the centrosome localization in naïve binucleated acinar cells. DAPI (magenta), pericentrin (cyan), annotation highlighting inter-nuclear space (yellow). Scale bar 5 μm. (D) Live-cell image series from H2B-mCherry mADOs showing mitotic binucleated cells. Top row: prior to mitosis, the nuclei orientation (green) is perpendicular to the cell’s long axis (yellow) which corresponds to the division axis (blue). Red annotations indicate the centrosome localizations with red arrows highlighting the microtubule orientation. Bottom row: the nuclei orientation equals the cell’s long axis and mitosis axis. H2B (white), nuclei annotation (magenta). Scale bar 5 μm. (E) Classification of the nuclei orientation either along the long or the short cell axis before multipolar mitoses based on live-cell imaging data from EGFP-Tuba mADOs. Grey line indicates a 50 % chance of alignment along the long axis representing a random nuclei orientation before mitosis. *P* values were calculated using a one-sided Student’s *t* test against the null hypothesis of equal distribution of nuclear orientations (50 %, n = 3 mice).

A higher likelihood of binucleated cells undergoing multipolar mitosis may be attributed to the restricted intracellular space resulting from metaplasia, which is associated with a significant decrease in cell size, combined with the increased spatial demands of two nuclei. We quantified the sizes of naïve acinar cells and metaplastic ADM cells from mADOs. As expected, cells that underwent ADM exhibit significantly reduced sizes (Figure. S2A-S2C). In addition, it has been proposed that the DNA itself might act as a physical barrier to prevent centrosomes from clustering to form bipolar spindle orientations in polyploid cells^50^. An unfavorable positioning of the centrosomes could further facilitate multipolar spindle arrangements. A previous study showed that centrosomes in binucleated G0/G1 cow trophoblasts cluster near the interspace between the two nuclei^51^. Such a positioning in combination with limited intracellular space might impair centrosome migration to the cell cortices during prophase. The positioning effect might even be stronger if the nuclei orient along the long axis of the cell, as it is the axis a cell generally tends to divide along (Hertwig’s rule)^52–54^. In addition, an orientation of the nuclei parallel to the axis of division, could prevent microtubule fibers from proper attachment to all kinetochores of both nuclei as the nuclei would shield each other in a bipolar spindle geometry. To resolve this issue and pass the spindle assembly checkpoint, a multipolar spindle orientation might be more favorable. Thus, we hypothesized that the cell size reduction induced by ADM, could force the nuclei in binucleated cells to orient along the long/division axis creating an unfavorable initial situation for bipolar mitoses. To test our hypothesis, we first stained naïve acinar cells for the centrosome marker pericentrin to assess centrosome positioning during G0/G1. In accordance with the data from cow trophoblasts, centrosomes in binucleated acinar cells cluster near the interface of the two nuclei (Figure 2C). We next assessed the nuclear orientation of binucleated cells just before mitosis using live cell imaging of EGFP-Tuba mADOs. We distinguished orientations along the short and the long/division axis of the cell and classified the subsequent mitosis as bipolar or multipolar (Figure 2D, Movies S3-5). Intriguingly, we found that multipolar spindle formations were always accompanied by nuclei orientations along the long/division axis of the cell (Figure 2E). In contrast, most bipolar divisions were performed with nuclei positioned along the short axis of the cell, perpendicular to the cell’s division axis. Of note, a fraction of binucleated cells exhibited nuclei orientations along the long axis, forming initial multipolar spindles but managed to resolve the multipolar geometry by centrosome clustering and still divided in a bipolar manner. These findings indicate that nuclei orientation and cell size have a pivotal impact on the spindle geometry and the number and ploidy of daughter cells.

Taken together, our findings define a new role for binucleated pancreatic acinar cells as facultative progenitors, that can transdifferentiate and divide at early stages of pancreas regeneration replenishing lost pancreatic tissue with two or more cells per mitotic passage, creating highly dynamic changes in nuclear numbers and cellular ploidy. This also implies that proliferating polyploid glandular cells are not unique to liver tissue. Instead, it highlights a conserved principle across multiple glandular organs, where injury-induced proliferation of otherwise quiescent polyploid secretory cells occurs as a shared regenerative mechanism. However, the simultaneous presence of metaplasia-induced cell size reduction increases the risk of multipolar mitosis, potentially contributing to tumor initiation over time.

### Polyploid mitosis leads to mitotic errors and micronuclei acquisition

Our next aim was to investigate, whether “scheduled polyploidy” as it occurs in binucleated acinar cells indeed represents a vulnerable state for chromatin segregation errors and CIN. Here, we use CIN broadly to encompass ongoing chromosome mis-segregation and its downstream genomic consequences, including micronuclei formation and structural variation. Studies of mammalian cell lines have demonstrated many times that polyploidy in its unscheduled form is perceived as a mutational process and thus is considered as a key step towards aneuploidy and the generation and evolution of cancer genomes^13,43^. A major contributor to the vulnerability of a polyploid cell to acquire CIN, lies in its tendency to form multipolar spindle geometries which possess an especially high chance for merotelic microtubule attachments and lagging chromosomes^13,55–61^. Thus, we acquired live-cell imaging data of H2B-mCherry mADOs and thoroughly screened mitotic events. We especially focused on early stages of mADO formation to capture the first binucleated cell divisions and acquire data as close as possible to an *in vivo* setting. We observed that the formation of mADOs is initiated by arranging cells from a primary acinus cell cluster into a spherical shape featuring a big lumen (Movie S6). This step lacks any cell divisions, in line with our previous BrdU assay, and recapitulates the formation of tubular complexes as seen in ADM in tissue. Strikingly, we find that the first binucleated cell divisions that occur after 24-48 h are frequently (∼47% ± 10.2%) accompanied by chromosome segregation errors such as lagging chromosomes and DNA anaphase bridges (Figure 3A, Movie S6). These errors eventually resulted in regressed cleavage furrows leading to failed cytokinesis and additional whole genome doubling of the single daughter cell (Figure S3A). Besides representing an obstacle for cleavage furrow ingression, lagging chromosomes might also end up forming micronuclei as they often are not integrated into the main nucleus. Since we frequently observed multipolar spindle formations as well as lagging chromosomes and chromatin bridges in polyploid acinar cells during ADM, we screened for micronuclei by immunofluorescence staining of wild type mADOs (Figure 3B). We detected micronuclei in almost all organoids with a peak of micronucleus abundance at 4 to 5 days of culture. This probably reflects the time when most of the binucleated cells in a forming mADO divided once.

**Figure 3.**
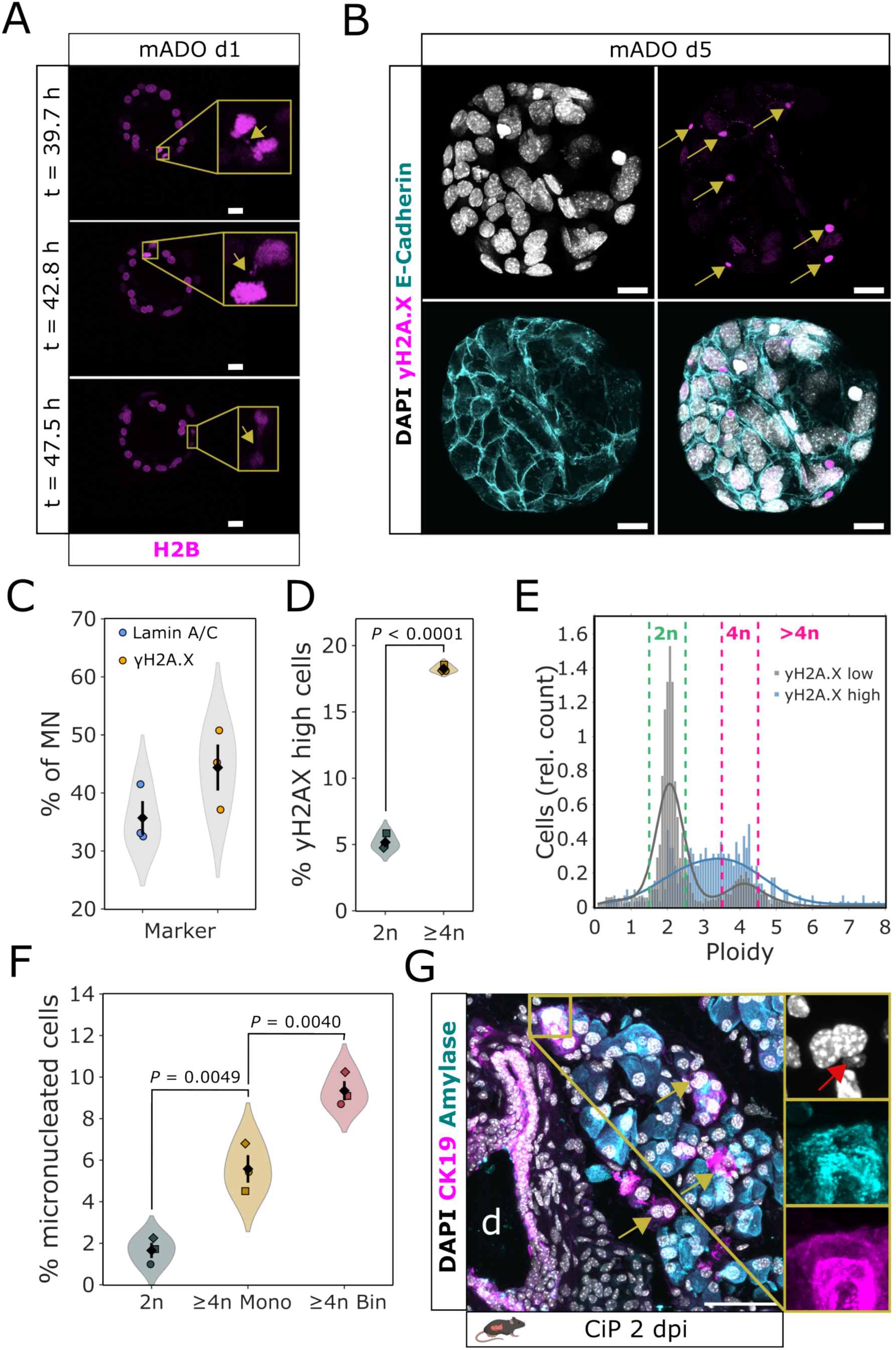
Polyploid acinar cells are predisposed to mitotic errors and micronuclei formation. (A) Live-cell image series from H2B-mCherry mice showing mitotic errors formed in proliferative binucleated cells at the onset of mADO formation (d1). H2B (magenta), yellow boxes indicate zoomed regions highlighting mitotic errors at different time points. Scale bar 20 μm. (B) IF staining from whole-mount d5 mADOs for γH2A.X (magenta) and E-Cadherin (cyan). Yellow arrows highlight micronuclei with strongly active DNA damage response in mADOs. Scale bar 20 μm. (C) Micronuclei characterization of d5 mADOs based on nuclear envelope composition (Lamin A/C low signal) or DNA damage response (γH2A.X high signal). Data is shown as percentage of all assessed micronuclei (n = 3 mice). (D) Violin plot showing γH2A.X intensity quantification in d5 mADOs comparing diploid (2N) and polyploid (4N) cells. *P* value was calculated using a two-sided paired Student’s *t* test (n = 3 mice). (E) Ploidy distributions by histograms and KDE curves of γH2A.X high and γH2A.X low cells in d5 mADOs (n = 3 mice). (F) Micronuclei abundance in d5 mADO based on percent of micronucleated cells according to ploidy and nuclear number: diploid (cyan), polyploid mononucleated (yellow), polyploid binucleated (grey). *P* values were calculated by one-way ANOVA followed by Tukey’s post hoc test (n = 3 mice). (G) IF image from mouse cerulein-induced pancreatitis at 2 dpi. Pancreatitis sections stained for α-Amylase (cyan) and CK19 (magenta). Yellow magnification box shows micronucleated acinar cell with red arrow indicating micronucleus. Yellow arrows indicate CK19-/Amylase-double-positive binucleated acinar cells, d: duct. Scale bar 50 μm.

Micronuclei are believed to play key roles in tumor initiation and evolution by promoting DNA damage and mutagenesis, which may give growth advantages to genetically unstable cells^62,63^. Their tumorigenic effect is linked to chromothripsis – the catastrophic shattering and reassembly of whole or partial chromosomes^21,64^. Micronuclei often lack nuclear membrane components, leading to instability and rupture, which contributes to the chromothripsis^64–68^. While these events are frequently observed in cancer cells, engineered or treated cultures, and tumors (with chromothripsis seen in about half of human cancers)^69,70^, they are rarely observed in normal cells, except in developmental contexts like germline and congenital disorders^71–73^. To assess whether MN envelope instability and chromothripsis occur during ADM, we stained mADO cells for the nuclear envelope component Lamin A/C and the DNA damage marker *γ*H2A.X (Figure 3B, Figure S3B). At d5, approximately 67% of all micronuclei stained negative for Lamin A/C, indicating differences in MN lamina integrity and composition and potential envelope instability in a majority of MN as compared to nuclear membrane (Figure 3C). Notably, 44% of the d5 micronuclei in mADOs exhibited massive DNA damage as indicated by strong *γ*H2A.X immunofluorescence signals, consistent with chromothripsis-like catastrophic DNA damage (Figure 3B, 3C)^64,74^. Since MN can be easily detected and quantified, as opposed to transient chromatin segregation errors, we further focused on the quantification of micronucleated cells and *γ*H2A.X signal intensities in mADO cells as well as determining their underlying DNA content. To this end, we developed an experimental and image analysis pipeline which includes the dissociation of mADOs into single cells, fixation onto microscopy slides via Cytospin centrifugation, staining for α-Tubulin, the cell cycle marker geminin and the DNA damage marker *γ*H2A.X as well as nuclear staining using DAPI (Figure 3C). Detailed image analysis workflow is described in the methods section. At d5 in culture, we found that the amount of *γ*H2A.X high cells is strongly increased in the polyploid fraction of mADO cells (Figure 3D). Additionally, our analysis revealed that the ploidy distribution of *γ*H2A.X high cells is generally shifted towards cells with higher DNA content, with many cells showing an intermediate ploidy, presumably indicating aneuploid chromosome sets (Figure 3E). In contrast, *γ*H2A.X low cells showed a more distinct bimodal distribution of diploid and tetraploid cells.

We further classified segmented cells at d5 by the presence of micronuclei based on the underlying DNA content and nuclear number. We observed the highest abundance of micronuclei in binucleated polyploid cells (9.37%). These results indicate that micronuclei formation often co-occurs with cleavage furrow regression leading to failed cytokinesis as described above. Our assay does not consider cells that underwent multipolar mitoses and ploidy reduction to become diploid while still being able to carry micronuclei generated in the ploidy reducing mitoses or earlier. Mononucleated polyploid cells still exhibited a greater proportion of micronucleation compared to mononucleated diploid cells (Figure 3F). Together, this implies that the true number of micronuclei linked to an initially polyploid genome could even be higher.

To validate our findings in an *in vivo* setting, we revisited the cerulein-induced pancreatitis model and stained tissue sections at 4 dpi for α-Amylase and CK19 to detect acinar cells that underwent ADM. We regularly found binucleated α-Amylase/CK19-double-positive cells carrying a micronucleus (Figure 3G). Of note, no micronuclei were detected in naïve freshly isolated acinar cells, indicating that those MN were formed during the inflammatory response upon the cerulein treatment.

Our results indicate that early cell divisions of polyploid transdifferentiating acinar cells are especially prone to undergo chromatin segregation errors eventually leading to micronuclei formation and chromothripsis-like DNA damage.

### Micronucleated acinar cells stay proliferative and acquire genomic structural variations

An ongoing question, that is regularly discussed in the light of tumor formation and evolution, addresses the fate of aneuploid/micronucleated cells. Excessive unrepaired DNA damage often leads to cell cycle arrest paired with increased autophagy which ultimately might lead to cell death and clearance^75,76^. Such protection can be normally achieved by cell-intrinsic pathways^77–80^. In contrast, it was also shown that micronucleated cells can re-enter cell cycle and pass their aneuploid genome to daughter cells^81^. Importantly, these findings were obtained in 2D cultured cells upon treatment with cytotoxic chemicals and/or ionizing radiation. However, to what extent micronucleated cells re-enter cell cycle and complete further mitoses under physiological conditions have not been previously examined. Using live-cell imaging of H2B-mCherry mADOs, we found that micronucleated cells are indeed capable of maintaining proliferative capacities (Figure 4A, Movie S7). In these cells, the micronucleus eventually gets reintegrated into the main nucleus and further mitotic defects can be commonly observed. Those reintegration events would be especially likely if the micronuclear DNA contains centromeric regions, which possess a certain degree of protection against DNA damage and can facilitate microtubule binding. To further delineate the full process of a mitotic error with micronuclei formation and subsequent proliferation, we subjected H2B-mCherry/mG mADOs to long-term live cell imaging over the course of 4 days, starting at d3 and using an inverted light-sheet fluorescence microscope. Tracking of the individual nuclei and cells based on the mCherry and GFP signals allowed us to reproduce the lineage relations in the growing organoid (Figure 4B). First, we noticed that the two daughter cells from an aberrant binucleated cell division behaved differently. While the MN acceptor cell remained proliferative (highlighted in red), the donating cells stopped proliferation, potentially because of the chromosome loss. Secondly, the proliferative MN acceptor cells did not show any growth disadvantages compared to other cells and had a similar cell cycle length. Generally, we measured an average doubling rate of 16.2 h. A similar experiment has been performed on mouse intestinal organoids, that acquired mitotic errors due to a highly proliferative phenotype with a rapid doubling rate of approximately 10 h^82^. While these cells potentially acquired mitotic errors because of their high (error prone) turnover rate, we propose that the tendency of pancreatic acinar cells to perform aberrant mitoses is inherently linked to the combination of polyploidy/binucleation and metaplasia.

**Figure 4.**
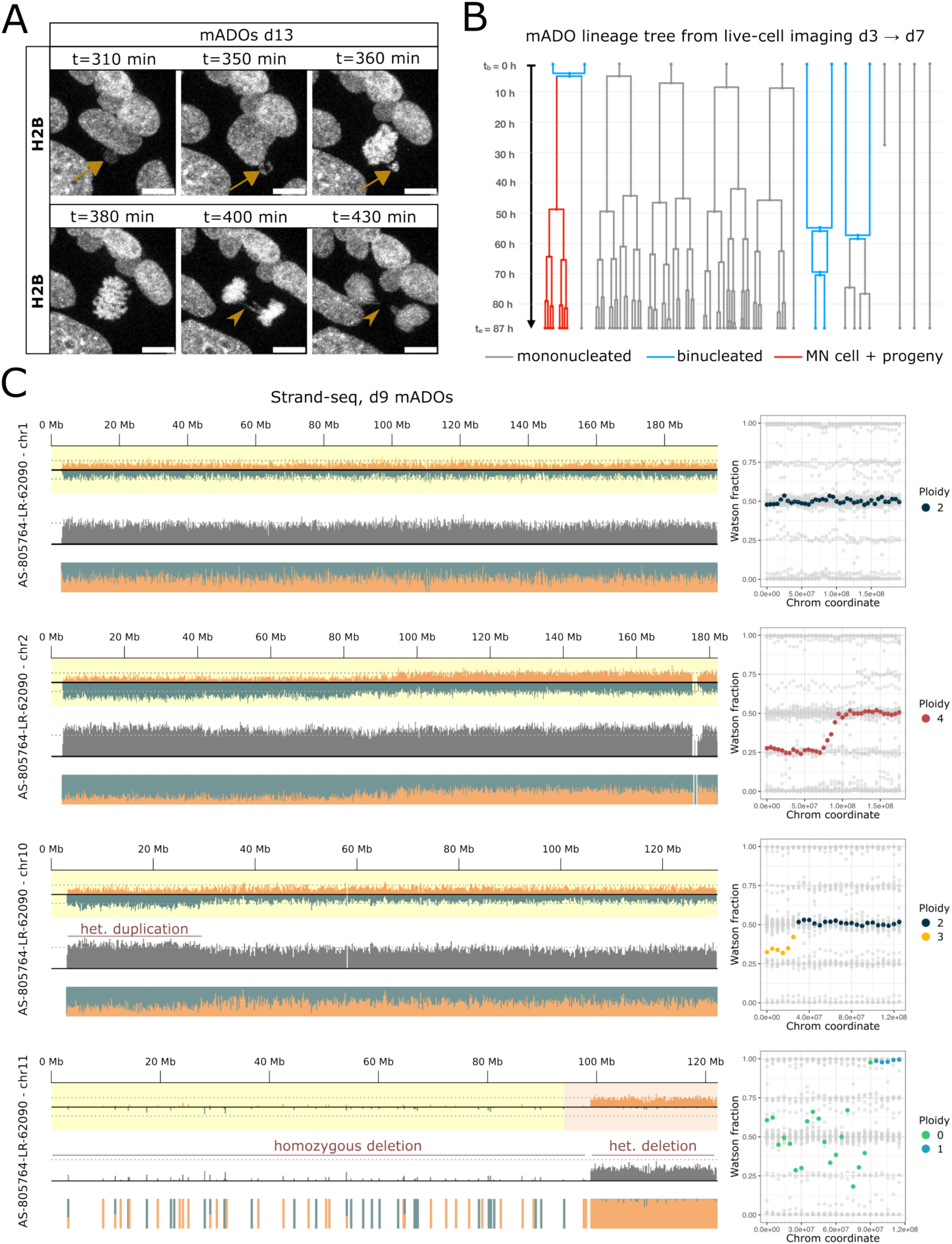
Micronucleated acinar cells remain proliferative and undergo repetitive segregation error cycles. (A) Live-cell image series of a d13 H2B-mCherry mADO. White arrow indicates micronuclei reintegration upon mitosis. White arrow heads indicate a subsequent chromatin bridge in the same cell. Scale bar 20 μm. (B) Reconstructed lineage tree of a growing organoid (d3 to d7). Grey tracks indicate mononucleated, blue tracks indicate binucleated cells. Red tracks indicate the lineage of a micronucleated cell from a binucleated mitosis with retained proliferative capacities. (C) Single-cell Strand-seq plot showing: a diploid chromosome (chr1), a tetraploid chromosome (chr2), a heterozygous duplication at the start of chr10, and a deletion in chr11. Left panels display coverage tracks with read depth divided by Watson and Crick strands, total read depths, and normalized Watson/Crick fractions, at a 200 kb bin resolution. The dotted lines represent the median value for the cell. Right panels show Watson fraction versus chromosomal coordinates, colored by ploidy, to highlight genomic gains or losses across chromosomes, in 5 Mb steps (smoothed over a 10 Mb window); gray points in the background represent data from the same chromosome in other cells.

To assess the genomic landscape of polyploid acinar cells after proliferation on the molecular level, we subjected cells from day 9 mADOs to single-cell whole genome sequencing. We employed Strand-seq, which gave us two advantages. First, by sequencing only the template strand of a single cell, haplotype information enables distinction of homologous chromosomes and detection of complex genomic rearrangements. Second, Strand-seq requires the processing of cells which have BrdU incorporated into the nascent strand from the previous replication phase. This allowed us to enrich for cells that have divided and potentially underwent chromatin segregation errors. To specifically sort for diploid and polyploid cells, we developed a two step-sorting strategy based on the fluorescent ubiquitylation-based cell cycle indicator (Fucci2) model (Figure S4A). This enabled us to specifically exclude S-, G2- and M-phase cells, which might possess an increased DNA content despite being diploid and sort cells based on ploidy. We sequenced 96 nuclei and obtained 42 high-quality genomes (22 from diploid and 20 from tetraploid nuclei). The average ploidy of each cell was assessed by visual inspection of Strand-seq results as shown in Figure S4B-D. We identified five cells exhibiting whole-chromosome aneuploidies and/or major structural variants (SVs). Among these, one cell displayed a complex ploidy structure characterized by some chromosomes gaining or losing one or two copies, or having segments entirely deleted, while other chromosomes remained intact and present in two or four copies (Figure 4C). Notably, the presence of extensive deletions in otherwise karyotypically normal cells suggests that the deleted regions may have originated from a lagging chromosome that was lost during mitosis (Figure 4C). Additional aneuploidies included the loss or gain of a full chromosome copy (in both diploid and tetraploid cells), leading to a pentaploid state, and triploid loci (Figure S4E).

Our findings demonstrate that organoids derived from primary naïve acinar cells acquire features typically found in tumors and that these transformations predominantly originate from polyploid cells. Mitotic errors and catastrophic DNA damage has not been linked to the natively occurring polyploidy found in acinar cells so far. Chromothripsis from mitotic errors has been proposed as a potential mechanism for tumor development in the pancreas and our data provides evidence that polyploid acinar cells are prone to these events^83^.

### Single-cell proteomics reveals two major acinar states defined by ploidy, STING–ATR axis usage, and CIN adaptation

To uncover signaling pathways allowing polyploid acinar-derived cells to adapt to chromosomal instability and even remain proliferative during ADM, we performed single-cell proteomics (SCP) on 367 cells from d5 mADO cells. Cells from three WT mice were sorted into 2n, 4n, and 8n fractions prior to SCP profiling. Our SCP workflow generated high-quality proteomes, with a median of ∼4,000 proteins per cell and a total of >6,100 unique proteins detected across the dataset (see Data S1 and Methods). UMAP embedding of the proteomic data revealed two dominant clusters in d5 mADOs (Figure 5A, Figure S5A). One contained predominantly 2n and 4n cells and exhibited high STING1 expression (STING1 high) and low ATR expression (Figure 5A). The second cluster was enriched for 8n polyploid cells displaying the inverse pattern: STING1 loss and high ATR. STING1 expression declined with increasing ploidy, whereas ATR scaled positively with ploidy and was highest in the STING-low cluster (Figure 5A, B). Correspondingly, DNA damage response (DDR) scores were strongly enriched in the STING1-low cluster (Figure S5B). We validated the timing of STING1 downregulation by immunofluorescence (IF) analysis of mADOs at days 5, 8, and 16. A significant downregulation of STING1 across all time points in 4n and 8n cells relative to 2n cells (Figure 5C, Figure S5C). This difference was even higher in cells with micronuclei (MN) than in MN-negative ones (Figure 5D). To address if loss of STING1 was indeed linked to ADM *in vivo*, we examined acinar cells within the naïve and injured pancreas. Surprisingly, in the naïve pancreas, STING1 was not suppressed, but instead showed, slight enrichment in 4n and 8n polyploid acinar cells (Figure 5e). This indicates that STING1 loss is not an intrinsic feature of polyploidy per se. Notably, in 5-dpi cerulein-induced acute pancreatitis (CiP), STING1 was selectively downregulated in polyploid acinar cells (Figure 5F), mirroring the ADM-associated suppression observed in organoids. These results suggest that STING1 suppression is an acquired adaptation triggered when polyploid acinar cells undergo ADM, re-enter the cell cycle and experience CIN, rather than an intrinsic property of polyploidy. STING1 downregulation has been proposed to represent a CIN-adapted state uncoupling tumor cells from interferon signaling in ovarian cancer exhibiting whole genome duplication^84–86^. To examine the role of STING1 in ADM, we treated freshly isolated acinar cells with the STING1 inhibitor H-151, which blocks STING1 palmitoylation, Golgi trafficking and degradation^87^. Further, single cell proteomics analysis of additional 367 H-151-treated d5 mADO cells showed that inhibited cells were distributed across both the STING1 high and STING1 low clusters, and additionally revealed a distinct cluster of polyploid STING1 high cells with low ATR and high KRAS (Figure 5G, Figure S5D). Although these cells expressed high levels of STING1, they were positioned closer in proteomic space to the STING1 low cluster than to the main STING1 high population. Within the STING1-high cluster, where H-151 is expected to act, the interferon response was reduced (Figure 5H). This is consistent with STING1 promoting interferon signalling, as interferon responses positively correlated with STING1 expression in untreated cells. Gene set enrichment analysis revealed that H-151 treatment induces multiple transcriptional and signalling changes in the STING-high cluster, whereas the STING-low cluster was largely unaffected (Figure S5E, S5F). Notably, the small subset of STING1 high/IFN low/KRAS high cells is consistent with a model in which STING1 inhibition dampens the interferon program and, in a subset of cells, permits secondary KRAS upregulation that enables escape from senescence or apoptosis and supports continued proliferation (Figure 5G). Conversely, the lack of a DNA damage response in these rescued cells suggests that STING1 downregulation may occur downstream of DNA damage to support adaptation and continued proliferation. Consistent with this, STING1 inhibited organoids exhibited a growth advantage compared to untreated controls, which manifested as an increased density of large organoid structures at d5 (Figure 5I, 5J). Together, our data show that adaptation to CIN via STING1 repression and DDR execution is not exclusive to full-blown tumors but occurs in polyploid regenerative cells in pre-cancerous settings, thus opening a critical window for potential malignant transformation.

**Figure 5.**
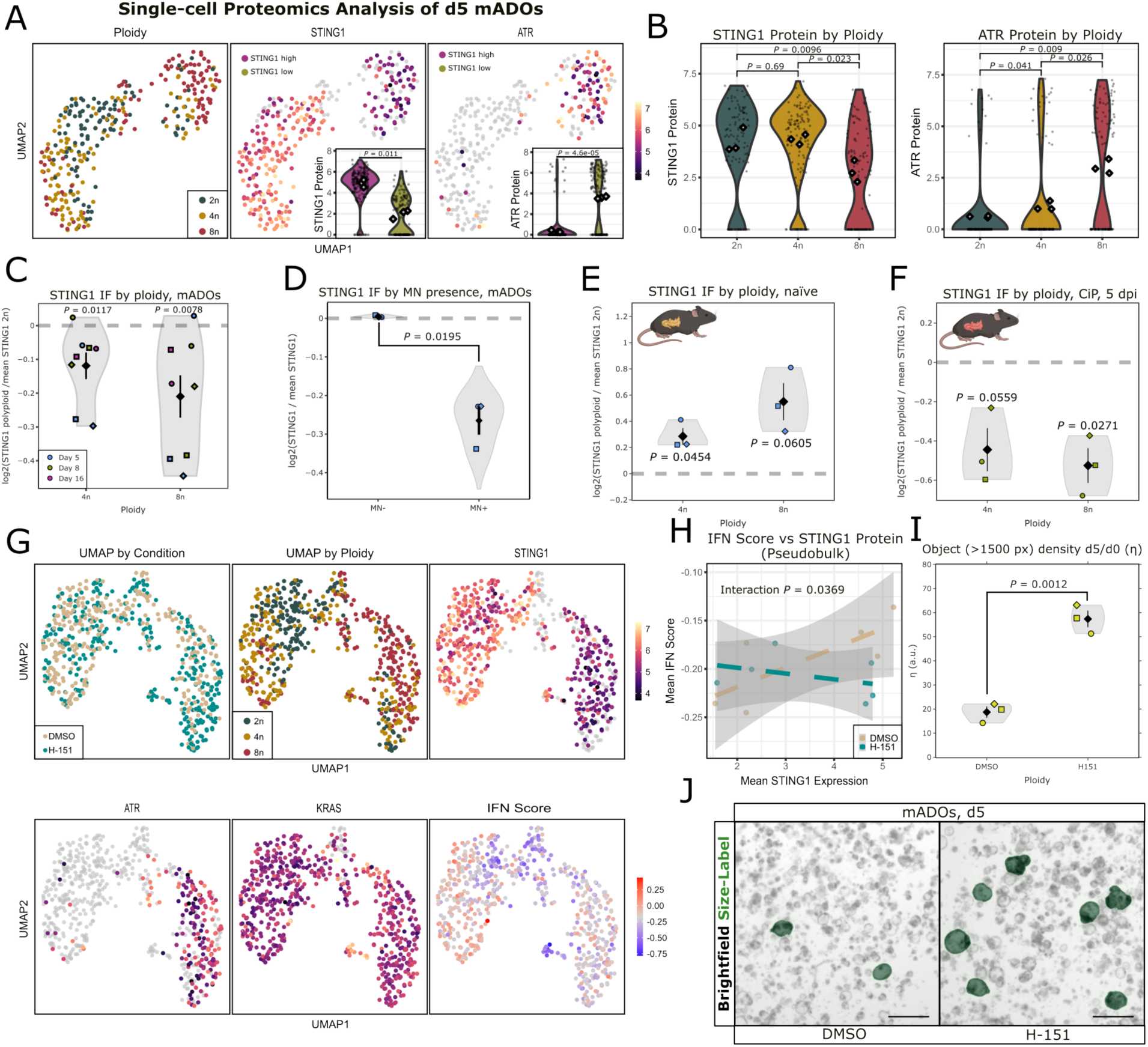
Polyploid acinar cells adopt a CIN tolerant state by repressing STING1 and upregulating ATR. (A) UMAP plots of single-cell proteomic data displaying ploidy labels (from FACS sorting) and STING1 and ATR protein intensities in untreated (DMSO) mADO cells; gray dots indicate cells in which the protein was not detected. Violin plots show the respective protein intensities between the two STING clusters (Figure S5A), with diamond shaped points indicating pseudobulk means (n = 3), on which a paired Student’s *t* test was computed. (B) STING1 and ATR protein expression stratified by ploidy. Pairwise *P* values were computed with paired Student’s *t* test and corrected by Holm-Bonferroni (n = 3 mice). (C) IF quantification of STING1 fluorescence signal intensities in d5, d8 and d16 mADOs. Intensities are represented for polyploid cell states as log2 fold changes relative to the corresponding median diploid (2n) STING1 intensities (dashed grey line). *P* values were calculated using a one-sided Student’s *t* test against the null hypothesis of a theoretical mean log2 fold change of 0 (n = 3 mice). (D) IF quantification of STING1 fluorescence signal intensities in d5 mADOs. Intensities are represented for micronucleated (MN+) and non-micronucleated (MN-) cells as log2 fold changes relative to the corresponding median (2n) STING1 of all cells (dashed grey line). *P* values were calculated using a paired Student’s *t* test (n = 3 mice). (E) IF quantification of STING1 fluorescence signal intensities in naïve primary mouse pancreatic acinar cells. Intensities are represented for polyploid cell states as log2 fold changes relative to the corresponding median diploid (2n) STING1 intensities (dashed grey line). *P* values were calculated using a one-sided Student’s *t* test against the null hypothesis of a theoretical mean log2 fold change of 0 (n = 3 mice). (F) IF quantification of STING1 fluorescence signal intensities in injured (CiP) mouse pancreatic acinar cells at 5 dpi. Intensities are represented for polyploid cell states as log2 fold changes relative to the corresponding median diploid (2n) STING1 intensities (dashed grey line). *P* values were calculated using a one-sided Student’s *t* test against the null hypothesis of a theoretical mean log2 fold change of 0 (n = 3 mice). (G) UMAP plots of single-cell proteomic data. H-151-treated and DMSO-treated ADO cells are annotated by experimental condition, ploidy labels, protein intensities (STING1, ATR, KRAS), and interferon (IFN) score. (H) Correlation between IFN score and mean STING1 expression calculated on pseudobulks (grouped by STING cluster, treatment, and mouse). The reported *P* value represents the interaction term between mean STING1 expression and condition, indicating that the inhibitor uncouples the IFN response from STING1 protein levels. (I) Densities ratios for large objects (>1500 px) in DMSO control and H-151 mADO cultures at d5. Object counts at d5 are normalized to the d0 overall object density of the same well. *P* value was calculated using a paired Student’s *t* test (n = 3 mice). (J) Representative brightfield images of d5 mADOs from DMSO control and H-151 treated samples. Green overlay highlights organoid structures that were segmented with areas larger than 1500 pixels (size threshold for quantification). Scale bar 20 μm.

### Polyploid cells in lactating mammary gland organoids acquire mitotic defects and micronuclei

While the pancreas maintains polyploid cells throughout the lifetime, the mammary glands in humans and mice only generate secretory polyploid cells during lactation when there is the need to nourish their offspring^8^. During pregnancy, luminal cells in the mammary gland differentiate into secretory alveolar cells capable of producing and secreting milk. After weaning, the mammary gland undergoes post-lactational involution, during which these alveolar cells either revert to a quiescent state, leading to a reduction in their size, or undergo apoptosis^88^. This makes the mammary gland a system of particular interest, because the non-lactating organ serves as an ideally suited control when studying naïve polyploidy. In addition, preclinical and epidemiological studies indicate that dysregulated post-lactational involution may contribute to tumor development^89^. Thus, to further generalize our findings from the pancreas to other secretory organs, we established lactating mouse mammary-derived organoids (mMDOs) with a near-physiologic cell composition of Keratin 5 expressing basal and Keratin 8 expressing luminal cells harboring a subpopulation of polyploid lactating (beta-Casein-positive) alveolar cells (Lummer, Brunken et al., *Pregnancy Hormone Treatment as a Tool to Induce Polyploidy in Mammary Gland Organoids*, 2026, under revision in STAR Protocols). The lactating capacities of these organoids rely on prolactin treatment after a morphogenesis phase, while the polyploidization is triggered by the addition of other known pregnancy-related factors such as progesterone, estrogen and EGF in an intermediate pseudo-pregnancy step (Figure S6A, S6B).

To assess the behavior of polyploid luminal cells, we performed confocal live-cell imaging of H2B-mCherry/mG mMDOs that have been subjected to hormone withdrawal thereby mimicking post-lactational involution^90^. Besides apoptotic cells, we noted that some binucleated cells remain proliferative after polyploidization and often divide via multipolar reductive mitosis (Movie S8). We further found that most of the cycling alveolar cells (beta-Casein-positive, geminin-positive) were polyploid and exhibited 6 times higher levels of *γ*H2A.X than their diploid counterparts (Figure S6C-S6E). We additionally detected micronucleated cells with retained proliferative capacities (Figure S6F). Taken together these results extend our findings from the exocrine pancreas to other glandular tissues that undergo tissue remodeling processes. We demonstrate that binucleated polyploid luminal cells of the mammary gland that emerge in response to pregnancy hormone treatment, represent a vulnerable state that might lead to mitotic defects and genomic instability. In addition, conditions like mastitis, which arises from blocked milk ducts or excess milk production during lactation could additionally activate proliferation of polyploid alveolar cells^91^. While high parity and extended breastfeeding were traditionally considered protective against breast cancer, postpartum-associated breast cancer challenges this notion, as nulliparous women can have a lower risk before age 40. The risk for parous women peaks five years postpartum and remains elevated for 24 years, with each year of delayed childbearing increasing breast cancer risk by 3.5%^92^. This may be partly explained by genomic instability in binucleated luminal cells, immune suppression, or mastitis during lactation.

## Discussion

### Polyploidy in naïve and regenerating glands

In glandular organs, differentiated epithelial cells can regain proliferative competence in response to injury or physiological demand, as seen during acinar cell–driven pancreas regeneration and during alveolar differentiation and involution in the mammary gland. Our study shows that polyploid cells in these organs do participate in this process.

We demonstrate a remarkable plasticity of binucleated polyploid acinar cells upon injury. While highly quiescent during homeostasis, binucleated acinar cells acquire the license to divide after injury by undergoing ADM, a key step in both, pancreas regeneration and tumorigenesis. After ADM, binucleated acinar cells exhibit a full cell cycle and can divide into two and eventually three cells leading to ploidy reduction and depletion of the binucleated fraction of acinar cells. During the post-regenerative phase, the number of binucleated acinar cells are restored by endomitosis. Endomitosis has also been proposed as the main mechanism for hepatocyte and acinar cell binucleation during development with potential implications in organ size control^93,94^. Thus, binucleation during regeneration could signal growth stop and resolution of the regenerative process.

The generation of more than 2 cells in a single round of cell division during pancreas regeneration stems from an increased number of centrosomes in polyploid cells and the resulting formation of more than 2 spindle poles during mitosis. Our results demonstrate that such multipolar spindle formations are more likely in binucleated than polyploid mononucleated acinar cells due to unfavorable centrosomal and nuclear positioning after metaplasia-induced cell size reduction. Cell culture experiments show that polyploid DLD-1 and RPE-1 cells quickly reduce their extra centrosomes, likely as an adaptive mechanism to prevent harmful chromatin segregation errors^44,47^. This process involves asymmetric clustering, where centrosomes unevenly distribute (e.g., one centrosome on one side and three on the other in tetraploid cells), eventually producing polyploid cells with only one centrosome. These single-centrosome polyploid cells may have a growth advantage over those with multiple centrosomes due to more accurate chromosome segregation, leading to a selective loss of cells with multiple centrosomes. Of note, organisms such as the allotetraploid *Xenopus laevis* have evolved similar mechanisms to ensure faithful chromatin segregation^95^. However, such adaption does not seem to occur during pancreas development, polyploidization and homeostasis, because binucleated acinar cells still possess extra centrosomes. We also demonstrated that the post-regenerative time is characterized by the re-binucleation of the acinar cell population via endomitosis which would lead to newly generated cells possessing more than one centrosome in G0/G1. A better understanding of ploidy and centrosome dynamics and their impact on the regenerative potential in glandular organs could lead to novel approaches to modulate and enhance tissue repair.

### Scheduled polyploidy as a potential origin of genomic instability

The polyploid state of a cell has been described controversially in relation to cancer. While many studies show detrimental effects of extra centrosomes, leading to chromatin segregation defects and DNA damage combined with higher resilience to undergo apoptosis, other studies propose a buffering effect of the extra copies of tumor suppressor genes, facilitating a certain degree of protection against loss of heterozygosity which has been primarily proposed for polyploid hepatocytes^13,96–99^. However, studies about tumor initiation often employ genetically engineered animal models or chemical treatments that would affect almost every cell in the corresponding tissues. For instance, diethylnitrosamine (DEN) mediated tumor induction leads to DNA alkylation adducts and tumorigenesis in hepatocytes^100^. This results in the formation of dozens of independent tumor nodules throughout the liver. Considering that DEN is a mutagen directly causing DNA damage, a buffering effect in polyploid cells after high dosage treatment seems reasonable. Further, though polyploid binucleated cells contribute to tissue regeneration, we show that diploid cells have a proliferative advantage at early stage of injury. However, “real-world” tumorigenesis is most often not induced by single high dosages of a mutagen affecting almost every cell in the organ as it is commonly performed in the lab but rather the results of multiple regeneration rounds following multiple injuries over many months and years, counting ageing as a mutagen itself. Regardless of whether tumor evolution is explained by a gradual or the more recently proposed punctuated equilibrium model, considerations of tumor origin should focus on cells that show the greatest tendency to DNA damage and genetic instability among the known risk factors^101^. Notta et al. proposed that, for the pancreas, the traditional view of mutation acquisition in a gradual manner does not match clinical observations of pancreatic cancer genomes per se^83^. Thus, they suggested that a significant fraction of PDACs arise through chromatin segregation error-related catastrophic events, like chromothripsis, which induce multiple mutations simultaneously. In agreement to this, 40% of PanIN lesions show chromothripsis^25^. Our data fits these observations very well, providing the binucleated polyploid state of an acinar cell as a potential origin for this punctuated equilibrium. Along the same line, epidemiological data link recurrent acute pancreatitis or chronic pancreatitis with acute flare-ups to a persistently higher long-term risk of pancreatic cancer compared to those with only a single acute episode or chronic pancreatitis without flare-ups^102^. We propose that while the organoid model replicates the acinar-to-ductal metaplasia (ADM) and proliferative phase characteristic of acute pancreatitis, the presence of EGF in the culture medium bypasses the requirement for acquiring a growth advantage, such as KRAS mutation, to drive tumor progression, thereby rendering the process irreversible. In contrast, in the acute cerulein model, although we observed aberrant mitosis of polyploid cells, the pancreas fully regenerated to normal within a year^7^, and only repetitive episodes lead to higher probability of having one of these aberrant cells gaining a proliferative advantage as it is observed in hereditary pancreatitis.

### STING suppression as an early adaptation to polyploidy-associated CIN

CIN is widely recognised as a potent trigger of cGAS-STING signalling through the formation and rupture of micronuclei, and sustained STING activity is generally considered cytotoxic or growth-limiting. Accordingly, several studies have shown that advanced CIN-high or WGD-high tumours downregulate components of the cGAS–STING pathway as a late immune-evasion strategy^84–86^. Our data extend this concept by demonstrating that STING1 suppression emerges much earlier, during the initial regenerative and pre-neoplastic phases of epithelial plasticity and not driven by exposure to the immune system. By contrast, polyploid acinar cells in vivo maintain, or even increase, STING1 expression. STING1 downregulation therefore represents an acquired adaptation that coincides with cell cycle re-entry, micronucleus formation and CIN, rather than a constitutive feature of polyploid identity. This suggests that despite high expression of STING in polyploid cells, acquired DNA damage leads to combined activation of ATR-dependent stress buffering and STING1 loss.

Several lines of evidence argue against the alternative interpretation that reduced STING1 levels simply reflect ongoing activation-induced protein turnover^103^. First, the STING1-low state persists across multiple time points and experimental conditions, consistent with a stable cellular programme rather than transient signalling dynamics. Second, STING-low cells remain viable and expandable despite elevated DNA damage, a phenotype incompatible with sustained STING activity^104^. Third, pharmacological STING1 inhibition with H-151, which blocks STING1 palmitoylation, Golgi trafficking and degradation, does not collapse the STING1-low cluster neither expands the STING-high cluster, indicating that activation-coupled turnover alone cannot account for reduced STING1 abundance. Together, these findings support a model in which early DNA damage creates selective pressure that favours STING loss as a CIN-adapted state, as shown by the anticorrelation between micronuclei presence and STING protein. Notably, STING inhibition also unmasks a small, otherwise non-permissive high-ploidy subpopulation characterised by high STING protein, KRAS and low ATR. This state is absent in untreated organoids, indicating that high STING activity induces death of polyploid cells, that can be rescued by STING inhibition. Consistent with this, STING1 inhibition is associated with the emergence of higher numbers of large organoid structures, suggesting that STING functions as an early surveillance barrier that shapes clonal selection during polyploid regeneration. Collectively, our results suggest that STING suppression is not merely a late tumour immune-evasion mechanism, but an early, cell-intrinsic adaptation that enables polyploid epithelial cells to tolerate CIN during tissue remodelling. This framework reconciles physiological polyploidy with tumour-associated CIN tolerance and implies that cancers arising from regenerating glandular tissues may inherit a pre-configured STING-low state. More broadly, our findings caution that therapeutic strategies aiming to exploit CIN-induced cGAS–STING activation, such as TREX1 inhibition, may be limited to tumors not exhibiting early adaptation to CIN in polyploid cell populations.

Together, our study uncovers a pivotal and previously unnoticed role of native and scheduled polyploidy especially multinucleation in genomic instability acquisition linked to regeneration in exocrine glands. We provide a new entry point to further study potential preventive, diagnostic and therapeutic strategies targeting the link between polyploidy and cancer genome evolution.

## Materials and methods

### Animal models

All animals used in this project were housed under specific pathogen-free conditions at a 12 h light dark cycle at 22 °C and fed *ad libitum.* All procedure and experiments were in accordance with the German Cancer Research Center (DKFZ) guidelines and approved by the Regierungspräsidium Karlsruhe. For this study, C57BL/6N mice are referred to as wild type (WT). The following transgenic mouse lines were used:

- H2B-mCherry (B6-Gt(ROSA)26Sortm3Sia): This mouse line constitutively expresses histone 2B (H2B) fused to mCherry, which endogenously labels cell nuclei.
- mG (B6-Gt(ROSA)26Sortm4.1(ACTB-EGFP)Luo/Amv): The mG construct specifically labels plasma membranes by utilizing a modified part of the membrane-bound domain of the MARCKS protein fused to GFP.
- EGFP-Tuba (B6-Gt(ROSA)26Sortm12.1Sia): These mice constitutively express EGFP fused to α-Tubulin, labeling microtubule fibers as part of the cytoskeleton as well as the mitotic spindle.
- Fucci2 (B6-Tg(Gt(ROSA)26Sor-Fucci2)#Sia): The Fucci2 mouse model expresses mCherry-hCdt1 in G1 and mVenus-hGeminin in S/G2/M phase.

H2B-mCherry and EGFP-Tuba mouse lines were kindly provided by Dr. Jan Ellenberg (EMBL Heidelberg) and the mG mouse line was kindly provided by Dr. Takashi Hiragi (former EMBL Heidelberg) and transferred to the DKFZ animal housings via embryo transfer. Fucci2 mice were a kind gift from Dr. Michael Milsom (DKFZ Heidelberg). Material transfer agreements with the original owners of the mouse lines were conducted.

R26-H2B-mCherry and mG mice were crossed to generate H2B-mCherry/mG double reporter mouse lines. All mice were aged 8-12 weeks when used for experiments.

### Cerulein-induced pancreatitis

Acute pancreatitis in mice was induced using the oligopeptide cerulein (Sigma-Aldrich, C9026) in WT mice. Cerulein was dissolved in phosphate-buffered saline (PBS) to a concentration of 5 μg/ml and administered hourly (50 μg/kg body weight) by 7 intraperitoneal (i.p.) injections on two consecutive days (14 injections in total). The day of the last injection was defined as 0 dpi. Three replicates of cerulein-treated mice were sacrificed and employed for pancreas extraction at 2 dpi, 4 dpi, 28 dpi and 91 dpi, respectively. To control for injection-induced injuries, three WT mice were i.p.-injected with comparable volumes of 0.9% NaCl and otherwise treated identically.

### Mouse pancreas extraction

To obtain pancreatic tissue, mice were anesthetized by an i.p. injection of 700 µl – 800 µl Ketavet (5.71 mg/ml)/Rompun (2.80 mg/ml) in 0.9% NaCl. Blood cells were cleared out by a left-ventricular perfusion using 20 ml Hank’s balanced salt solution (HBSS). In case of tissue fixation for cryo-sectioning, perfusion was continued with another 20 ml of 4% Roti-Histofix (Carl Roth). The pancreas was exposed by opening the abdominal cavity via a mid-abdominal vertical incision. Pancreatic tissue was extracted and attached adipose tissue was removed, if necessary. Until further usage, pancreatic tissue was kept in PBS on ice.

### Mouse acinar-derived organoid culture

The following solutions were prepared and sterile-filtered: S (4% bovine serum albumin (BSA) in PBS), R (1% BSA in PBS), D (1 mg/ml Collagenase IV in 0.25% BSA in PBS) and W (2% Penicillin-Streptomycin in PBS). Pancreatic tissue was extracted and collected in 10 ml solution W on ice. The tissue was separated from remaining fat and rinsed once more in 10 ml solution W. The tissue was then chopped into small pieces with a volume of approximately 1 mm^3^. Tissue pieces were collected, rinsed in 10 ml solution W and then digested in solution D for 30 min at 37 °C and 5% CO2. Every 5 min, the suspension was pipetted up and down using serological pipettes (5 ml) to further facilitate dissociation. The digestion product was filtered through a 100 µm cell strainer to remove pancreatic islets and the cell strainer was rinsed with 10 ml Solution R. Four centrifugation tubes (15 ml) were prepared, each containing 6 ml of Solution S. For each tube, 5 ml of the filtered cell suspension were gently transferred on top to achieve a BSA gradient by layer separation. Acini were isolated by a single centrifugation step at 50 x g and 4 °C for 2 min. The supernatants were, removed and the pellets were resuspended and washed with Solutions S and W, successively (50 x g, 4 °C for 2 min). After the last washing step, the pellets were resuspended in 500 µl Pancreatic organoid culture (POC) medium (1:1 DMEM/F12 + GlutaMAX^TM^ 2% (v/v) B27 serum free supplement, 1% (v/v) N2 supplement, 1% penicillin-streptomycin, 2 mM L-glutamine 20 ng/ml rhEGF, 20 ng/ml rhFGF2) each and pooled. At this point, acinar cells build clusters of approximately 4-10 cells per cluster. Since cell-cell contacts facilitate organoid formation for this cell type, no further dissociation steps were performed. Acinar cells were counted and mixed with ice cold unpolymerized Matrigel® according to a density of 250 cells/µl. For 3D organoid culture, a 20 µl-droplet of this mixture was placed in each well of the respective cell culture dishes and incubated for 20 min at 37 °C and 5% CO2 to facilitate Matrigel® polymerization, before POC medium was added. Acinar cells were cultured at 37 °C and 5% CO2 in either 24-well plastic plates for fixed-cell imaging experiments and Strand-seq, 10-well CELLview^TM^ Slides (Greiner Bio-One) for confocal live-cell imaging or Luxendo TruLife dishes for light sheet live cell imaging. The time point of cell seeding in Matrigel® was defined as day 0 (d0).

### Mouse mammary gland extraction

Female animals were euthanized via cervical dislocation. Following sacrifice, the mice were pinned in supine position, and the ventral area was disinfected with 70% ethanol. A nick was cut above the pubis with small surgical scissors and continued towards the thoracic nipples. Then, two lateral incisions were made, and the skin was peeled to reveal the inguinal mammary fat pads. For thoracic pads, the same procedure was repeated in the thoracic area. To remove the fat pads, they were squeezed and pulled with the forceps while making small incisions to remove them from the skin. Lastly, collected mammary glands were washed in 1x cold PBS and transferred to a 50 mL tube containing 1x Dulbeccos Modified Eagle Medium/Nutrient Mixture F-12 (DMEM/F12 (1:1) + GlutaMAX™) at room temperature (RT).

### Mouse mammary gland derived organoid culture

The generation of mouse mammary gland-derived organoids is based on existing protocols^90^ (Lummer, Brunken et al., *Pregnancy Hormone Treatment as a Tool to Induce Polyploidy in Mammary Gland Organoids*, 2026, under revision in STAR Protocols). After extraction, glands were quickly rinsed in 1x DPBS and transferred to a 100 mm petri dish for mincing with 2 sterile disposable scalpels. Then, minced mammary tissue was transferred to a 50 mL falcon containing 10 mL of pre-warmed digestion medium I (2 mg/ml Collagenase A, 0.25 % trypsin, 5 % fetal bovine serum (FBS), 10 μg/ml gentamicin in 1:1 DMEM/F12 + GlutaMAX^TM^) and incubated on an orbital shaker at 200 rpm at 37 °C for 40 min. After digestion, the tube was briefly vortexed and the content was transferred to a 15 mL falcon, which was centrifuged at 450 x g for 10 min. Resulting fat layer contains trapped mammary cells inside, so together with the supernatant it was taken to a new falcon and pipetted up-down to release the epithelial cells from the fat. The fat mixture was centrifuged again, and the pellet was combined with the main pellet to maximize the yield. The combined pellet was resuspended in 5 mL wash medium I (5 % FBS in 1:1 DMEM/F12 + GlutaMAX^TM^) and inverted 5-6 times, followed by centrifugation for 5 min at 450 x g. The supernatant was discarded and the pellet was resuspended with red blood cell lysis buffer (RBC) and incubated for 1 min at RT to get rid of red blood cells. After the incubation, 8 mL PBS/1%FBS was added into the tube, and it was centrifuged for 2 min at 200 x g. The supernatant was aspirated and the pellet was resuspended with 4 mL pre-warmed digestion medium II (5 mg/ml Dispase 2, 0.5 mg/ml DNAse I in 1:1 DMEM/F12 + GlutaMAX^TM^). The cells were incubated in digestion medium II for 5 min in a 37 °C water bath. 6 mL wash medium II (0.1 % FBS in PBS) was added to stop the reaction. Then the tube was centrifuged at 450 x g for 5 min, and the pellet was resuspended in 1 mL Morphogenesis Organoid Medium (MOM, 1:1 DMEM/F12 + GlutaMAX^TM^ supplemented with 1 % (v/v) ITS-X, 2 %, 1 % penicillin-streptomycin, 45 ng/ml rhFGF2). Freshly isolated primary mammary epithelial cells were mixed with growth factor reduced Matrigel® and plated in domes in 8 µL Matrigel® pre-coated 24-well culture plates for fixed-cell experiments or 10-well CELLview^TM^ Slides (Greiner Bio-One) for live-cell imaging (one dome per well, 20 µL of Matrigel® per dome, ∼1500 organoids per dome) to increase the stability of the domes. After the Matrigel® polymerization in upside down position for 35 min at 37 °C, the 3D organoid cultures were covered with pre-warmed MOM and incubated at 37 °C in 5% CO2 incubator. Fresh media change was done once in every 2 days periodically; all media were prepared freshly every time. For the experiments, the organoids were cultured in MOM for 6 days, followed by either another 6 days in MOM, 6 days in Pregnancy-Lactation Medium (PLM, MOM lacking rhFGF2 but supplemented with 120 ng/ml murine recombinant prolactin, 25 ng/ml rhEGF, 40 ng/ml β-Estradiol (E2), 120 ng/ml Progesterone (P4) or 6 days in PLM lacking β-Estradiol (PLM/-E2).

### Fixation and single-cell dissociation for mADOs and mMDOs

At given time points, mADO and mMDO cultures were dissociated using TrypLE for 20-30 min with visual validation of dissociation under the microscope. After dissociation, single cells were washed once in PBS and fixed for 15 min in 4% PFA for further usage.

### *In vivo* BrdU-assay

BrdU (15 mg/ml in 0.9% NaCl) was administered i.p. according to 150 mg/kg body weight to three C57BL/6N WT mice at 1 dpi after cerulein-induced pancreatitis. Mice were sacrificed and pancreas tissue was extracted at 91 dpi.

### *In vitro* BrdU-assay

mADOs from one C57BL/6N WT mouse were cultured in 24-well plates as described above. Starting at 0 h after cell plating, BrdU was added to POC medium to a final concentration of 10 μM for three wells. After 2 h incubation the cells from these three wells were dissociated and fixed and BrdU was added to another three wells. This procedure was followed 24 h consecutively.

### Immunohistochemistry (IHC) staining

After extraction, pancreatic tissue was post-fixed in 4% PFA for 2 h at 4 ◦C. For tissue dehydration, the pancreas was transferred into 30% sucrose in PBS and incubated at 4 ◦C until it settled on the bottom of the vial. The tissue was embedded in Tissue-Tek® O.C.T.^TM^ Compound and stored at −80◦C. Prior to antibody staining, the tissue was cut into sections of 20 µm thickness using a cryomicrotome (−20 °C) and applied to microscope slides (Superfrost®Plus, Thermo Fisher). The tissue was incubated in PBS at RT until the surrounding Tissue-Tek® O.C.T.^TM^ Compound was completely removed. Tissue sections were bordered using a hydrophobic pen. In case of BrdU staining, tissue sections were incubated in 2 M HCl for 30 min at 37 °C followed by neutralization with 0.15 M sodium borate buffer for 30 min at RT and three times washing in PBS for 5 min at RT. For permeabilization, the tissue was incubated in 0.1% Triton^TM^-X 100 in PBS for 30 min at RT. Blocking was performed by incubation in wash/block buffer (0.1% Triton^TM^-X 100 in PBS) for 1 h at RT. Primary antibodies were diluted in wash/block buffer according to Table S1 and applied to the tissue sections for primary staining over night at 4 °C. Afterwards, the tissue was washed three times with wash/block buffer for 5 min at RT. Secondary antibodies were diluted in wash/block buffer according to Table S1. Tissue sections were incubated in secondary antibody solution for 1 h at RT. After washing three times in wash/block buffer for 5 min at RT, the tissue was mounted using Fluoromount G, with DAPI (4’,6’-diamidino-2-phenylindole) and stored at 4 °C until data acquisition. Pancreatic tissue sections of human patients were provided by the tissue bank of the National Center for Tumor Diseases (NCT) and stained by the Institute for Pathology of the Heidelberg University Hospital. Human tissue sections were stained for E-cadherin with a nuclear using hematoxylin. All steps were performed according to the manufacturer’s protocol (VENTANA anti-E-cadherin (36) Mouse Monoclonal Primary Antibody, Roche Diagnostics).

### Immunocytochemistry (ICC) staining

After dissociation and fixation at given time points, single mADO and mMDO cells were fixed on microscopy slides by Cytospin^TM^ centrifugation (10 min at 8000 rpm). Cell areas were bordered using a hydrophobic pen. For permeabilization, cells were incubated in 0.1% Triton^TM^-X 100 in PBS for 30 min at RT. Blocking was performed by incubation in wash/block buffer for 1 h at RT. Primary antibodies were diluted in wash/block buffer according to Table S1 and applied to the tissue sections for primary staining over night at 4 °C. Afterwards, the cells were washed three times with wash/block buffer for 5 min at RT. Secondary antibodies were diluted in wash/block buffer according to Table S1. Cells were incubated in secondary antibody solution for 1 h at RT. After washing three times in wash/block buffer for 5 min at RT, cells were mounted using Fluoromount G, with DAPI (4’,6’-diamidino-2-phenylindole) and stored at 4 °C until data acquisition.

### Fixed-sample confocal imaging

IHC/ICC-stained tissue sections and cells were imaged as xyz stacks using a Leica TCS SP8 confocal microscope. The xy resolution was set to 1024 × 1024 pixels at a line frequency of 200 Hz (tissue sections) or 400 Hz (Cytospin^TM^ slides) and a z-step size of 2 µm (Δz = 2 µm). Tuneable spectral photomultiplier tubes were used as detectors. Images were acquired using a 40x immersion oil-based objective (Leica 40x Plan Apo NA 1.30).

### Live-cell imaging

Time-lapse imaging of mADOs and mMDOs was performed using a Nikon A1R confocal microscope as well as a Luxendo InVi SPIM light sheet microscope (long-term imaging for lineage tree reconstruction of mADOs only). Live-cell imaging using a Nikon A1R confocal microscope was performed with a 20x air objective (Nikon 20x Plan Apo λ NA 0.75) at z-step sizes of Δz = 2 µm. The xy resolution was set to 1024 × 1024 pixels at a line frequency of 200 Hz. Light sheet imaging using a Luxendo InVi SPIM was performed with a 10x illumination objective (10x CFI Plan Fl NA 0.3) and a 25x detection objective (Nikon 25x CFI Apo NA 1.1) at Δz = 1 µm and 2048 x 2048 pixels (Δx = 204 nm, Δy = 204 nm). The light sheet diameter was set to 1.7 µm.

Double-fluorescent H2B-mCherry/mG mice were used to detect single cells within organoids and to determine the number of nuclei per cell as well as NEB to anaphase durations. EGFP-Tuba mice were used to visualize spindle formations in mADOs to quantify and characterize multipolar division events. Time periods of imaging ranged from 8 h to 84 h depending on the experiment at a temporal resolution of 6 frames per hour (Δt = 10 min).

### Manual quantitative image analysis

Manual cell counting was performed for all experiments involving the analysis of tissue sections using the built-in Cellcounter tool in Fiji ImageJ (v2.14.0/1.54f). This includes the quantification of mononucleated and binucleated cells as well as CK19-, pHH3- and DCLK1-positive. Additionally, the quantification of amylase-expressing cells in mADOs was performed likewise. Manual image annotations for visualization purposes were drawn using Napari (v0.4.18) or Inkscape (v1.3.2). NEB to anaphase durations were determined by visual frame-to-frame inspection and quantification of time lapse movies of H2B-mCherry/mG mADOs using Fiji ImageJ (v2.14.0/1.54f). Bipolar and multipolar divisions in mADOs were assessed by screening EGFP-Tuba time-lapse movies for the corresponding type of division using Fiji ImageJ (v2.14.0/1.54f). The nuclear orientation axis of binucleated cells before bipolar or multipolar division was assessed by manually annotating and measuring the cell’s long and short axes in the last time frame before NEB becomes visible using Fiji ImageJ (v2.14.0/1.54f).

For the lineage tree reconstruction from long-term live-cell imaging experiment, raw data was cropped using the Fiji plugin BigDataProcessor2 (v1.7.1). Cropped images were subjected to denoising via Noise2Void (v0.3.3, https://github.com/juglab/n2v) and deconvolution via flowdec (v1.1.0, https://github.com/hammerlab/flowdec). After conversion to BigDataViewer (v6.3.0) xml/h5 files, the lineage tree was reconstructed using Mastodon (v1.0.0-beta-30). Track lengths were directly extracted from reconstructed lineage trees. Only full cell cycles were used to determine the average cell cycle length, starting at the F1 generation.

### Automated quantitative image analysis

Dissociated organoids were fixed and adhered to microscopy slides using Cytospin and cells were subsequently stained for α-tubulin, geminin, γH2A.X, and nuclei using DAPI. Alpha-tubulin or CK19 staining enabled the segmentation of cell bodies, while geminin was used to identify cells in G2/M (exhibiting doubled DNA content) or Ki67 for cycling cells for exclusion from further analysis. γH2A.X was used to detect DNA damage response, and DAPI staining provided a measure of DNA content. Confocal images were exported as both maximum and sum intensity projections for each of the four channels using Fiji/ImageJ (v2.14.0/1.54f). The maximum projection of the alpha-tubulin channel was employed for instance segmentation with Cellpose (v2.2.3), whose “Cyto 2” model was fine-tuned on a subset of α-tubulin images. For each segmented object, features including area and perimeter (derived from the masked labels) and the median and integrated pixel intensities for DAPI, γH2A.X, and geminin (obtained from the masked sum intensity projections) were extracted using numpy (v1.24.3) and scikit-image (v0.21.0). Corrected total cellular fluorescence (CTCF) and corrected mean cellular fluorescence (CMnCF) values were then calculated for each channel by subtracting the background fluorescence intensity 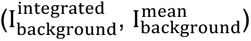 value multiplied with area of each object (A) from the object’s mean and integrated fluorescence intensities 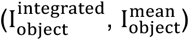 respectively:

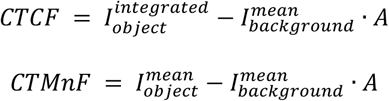

Each objects circularity was calculated using the objects area and perimeter (P) according to:

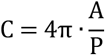

Area and circularity were used to flag mis-segmented objects (e.g., doublets or cells at image borders). Subsequent analyses focused on DAPI CTCF and CMnCF in the cases of geminin, γH2A.X or STING1, with the corresponding histograms smoothed via a Gaussian smoothing algorithm. For each channel, the largest peak was detected (local maximum approach via scipy.signal.find_peaks (v1.10.1)) and defined as diploid (2n) for DAPI, G0/G1 for geminin, and γH2A.X low for γH2A.X – an assumption verified by visual inspection. After Gaussian fitting around the largest peak with the peak center as an initial guess for the mean, the histograms were normalized according to the fitted mean of the normal distribution. In case of DNA content via DAPI CTCF, the values were additionally multiplied by the factor 2 to enable readability as ploidy values. Normalized fluorescence thresholds were established as follows: DAPI CTCF values of 1.5–2.5 (diploid), 3.5–4.5 (tetraploid), and >4.5 (higher polyploidy). For the normalized geminin and γH2A.X values, the thresholds were selected by plotting the corresponding CMnCF fluorescence intensity values against the normalized DAPI CTCF. To assess micronuclei presence and determine nuclear number, the maximum projection of the DAPI channel was cropped for each cell, based on the α-tubulin segmentation label, generating individual 90×90 pixels-sized cell patches. A classifier built on an ImageNet pre-trained EfficientNet B1 convolutional neural network (CNN) using Tensorflow (v2.13.0) was re-trained for 200 epochs on 1985 labeled images with a validation split of 0.2 and at a learning rate of 10^−5^ using an Nvidia RTX 4090 GPU, to categorize each cell patch into one of 5 classes (“mononucleated”, “binucleated”, “mononucleated micronucleated”, “binucleated micronucleated”, and “no cell” - representing mis-segmentation or debris):

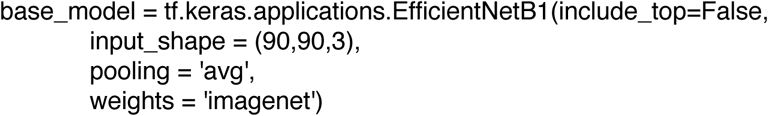

The training data was augmented using a random flip in horizontal and vertical orientations, random rotation as well as random contrast. The batch size was set to 16. The last 24 layers were additionally unfrozen to enable feature leaning from high-level features. The top of the architecture was complemented by adding a flatten layer, a dense layer with 500 neuron (ReLu activation) and another dense layer with 5 neurons (Softmax activation) for class output. The final model consisted of 7233676 parameters. For the loss function, Sparse categorical cross entropy was used. The model performance was tracked by validation loss and best weights were saved. The model performance was tested on test set consisting of 396 images from all 5 classes. The model performance was assessed by precision (positive predictive value), recall (sensitivity) and F1-score (harmonic mean of precision and recall). The resulting cell classification was integrated with the extracted features for downstream analysis. Full code for the image analysis pipeline is available upon request. In case of changes to the repository, all source code is of course also available upon request.

All classification results that went into the final quantifications were assessed by visual inspection of the classified image patches and manually corrected if necessary.

### Fluorescence-activated cell sorting (FACS) for Strand-seq

mADOs generated from Fucci2 mice were cultured in POC medium as described above. BrdU was added to the medium at 40 μM and incubated for 40 h. At d9, the cells were dissociated into single cells as described above. An initial sorting step was performed with intact cells to exclude G2 (mVenus^+^) cells and enrich for G0/G1 (mCherry^+^). At this, single cells were sorted into nuclear sorting buffer B containing Hoechst33258 and propidium iodide (PI) as described by the author of the Strand-seq protocol ^105^. After cell lysis, nuclei were sorted by Hoechst33258 signal to detect BrdU incorporation and PI signal to distinguish different DNA contents of nuclei (see Figure S4A). For further details regarding the selection process of sorting gates, we refer to the original Strand-seq protocol ^105^. Single BrdU-containing nuclei were sorted into 96-well plates containing 5 μl freeze buffer (42.5% (v/v) 2x Profreeze-CDM, 7.5% DMSO (v/v), 50% (v/v) PBS) and stored at −80 °C until further processing.

### Strand-seq

Strand-seq was used to detect structural variants in mADO cultures. The detailed protocol for Strand-seq library preparation is described in^105^. Sequencing was performed using a NextSeq 550 sequencing system (NextSeq 500/550 Mid Output Kit v2.5 (150 cycles)).

### Processing and analysis of Strand-seq data

Raw sequencing files were processed using the MosaiCatcher pipeline (v2.3.5)^106^ to align reads on mm10 genome, filter low quality cells, and obtain raw read counts at 200kb genomic bins. The strandtools package (https://git.embl.de/cosenza/strandtools) was then used to normalize binned counts and to generate, in addition to custom python code, coverage track plots.

For each genomic bin, the number of reads aligned to each strand defined the Watson Fraction (WF), that is, the number of reads aligned to the Watson strand over the total number of reads (sequencing depth) for each bin. For the ploidy of each nucleus, the presence of SVs was assessed by visually inspecting WFs and normalized total counts across each chromosome in each high-quality nucleus.

### mADO Culture and FACS sorting for single-cell proteomics

mADOs from three WT mice were cultured as described above. One half of the cells was incubated in POC medium (+0.008 % DMSO) and the other half in POC medium supplemented with 4 μM H-151 STING inhibitor (in DMSO). Medium was changed on d3, continuing H-151/DMSO treatment. At d5 mADOs were dissociated into single cells using TrypLE digestion and sorted using a BD FACSDiscover^TM^ S8 cell sorter. To determine DNA content levels, Hoechst333442 (15 μg/ml) was used in combination with with Reserpine (5 μM) to prevent Hoechst33342 efflux. SYTOX^TM^ Green was used for dead cell exclusion. Single cells were sorted into 384 well plates and immediately processed for single-cell proteomics analysis.

### Single Cell Proteomics LC–MS analysis

Peptides were analysed on a Vanquish Neo UHPLC system (Thermo Fisher Scientific) coupled to an Orbitrap Astral mass spectrometer (Thermo Fisher Scientific). Single cell proteomics data were acquired using Tune software (version 0.4 Thermo Fisher Scientific). Peptide separation was performed on an Aurora Rapid 8 cm nanoflow UHPLC column with integrated emitter (IonOpticks) operated in direct-injection mode at 50 °C, using an 18-min gradient in an 80 samples-per-day method. The mass spectrometer was equipped with a FAIMS Pro Duo interface and operated with an EASY-Spray source (both Thermo Fisher Scientific). A compensation voltage of −46 V was applied in FAIMS, and an electrospray voltage of 1.85 kV was used for ionization. MS1 survey scans were acquired in the Orbitrap analyzer at a resolution of 240,000 over an m/z range of 400–800, with an automatic gain control (AGC) target of 500% and a maximum injection time of 100 ms. MS2 spectra were acquired in data-independent acquisition (DIA) mode in the Astral analyzer using non-overlapping isolation windows of 20 m/z to cover m/z range of 400–800. The precursor accumulation time was 40 ms and the AGC target for MS2 scans was set to 800%.

### Single Cell Proteomics Data analysis

Raw data were analysed using Spectronaut (version 19.1.240806.62635, Biognosys). Searches were performed in DirectDIA+ mode with match-between-runs enabled, with each single cell as biological replicate in the search settings. Enzyme specificity was set to trypsin, and the default minimum peptide length was seven amino acids. Quantification was performed at the MS1 level. Carbamidomethylation of cysteines as a static modification was disabled for single-cell searches, as no alkylation step was included in the sample preparation. Data were searched directly against a mouse reference proteome (UniProt proteome UP000000589, reviewed, 17,240 protein entries; downloaded 12 April 2025) supplemented with an additional spectral library, and against the CRAPome contaminant database (118 protein entries). Single-cell protein intensity data were processed using Scanpy (v1.11.0). Quality control was performed by removing cells with fewer than 1,000 detected proteins or a total protein intensity sum exceeding 2.5×10^6, resulting in 672 high-quality cells. Missing values in the intensity matrix were zero-filled prior to log2-normalization. Dimensionality reduction was performed via Principal Component Analysis (PCA), followed by UMAP, using the first 20 principal components (PCs). A cluster identified as pancreatic stellate cells (PSCs) based on marker expression was removed, retaining 618 ADM cells. Following PSC removal, PCA was recomputed, and the dataset was integrated by plate using the harmony_integrate() function. UMAP embeddings were subsequently generated based on the first 30 PCs. For the DMSO-only sub-analysis (n=317), H-151 treated cells were excluded, and the dimensionality reduction steps were repeated. DNA damage repair scores (based on Tp53, Atm, Atr, Cdkn1a, and Brca1 intensities) and interferon scores (based on the gene list by Wu et al.) were calculated using the score_genes() function. Gene Set Enrichment Analysis (GSEA) was conducted using the gseapy Python package (v1.1.10) with the Reactome_Pathways_2024 pathway set, ranking genes by log2FC. Statistical analyses and plotting were performed in R (v4.5.0). Protein intensities between pseudobulks and score values were compared using paired t-tests, with Holm-Bonferroni correction applied where necessary.

### Statistical analysis and plotting for imaging data analysis

Statistical analyses were performed using scipy (v1.10.1). *P* values less that 0.05 are considered as statistically significant. Comparisons between two groups were performed using a paired Student’s *t* test in case of paired biological replicates or an unpaired Student’s *t* test in case of unpaired biological replicates. Comparisons between more than two groups were performed using analysis of variance (ANOVA) followed by a Tukey’s post hoc test. One-sided tests were used when a directional effect was hypothesized a priori; all other comparisons used two-sided tests. Graph visualization was done using Plotly (v5.17.0). Violin plots represent individual biological replicates as individual dots, in case of paired data with different symbols per replicate. Additionally, the mean is shown as black diamond and s.e.m. as black bars around the mean. Figures were prepared using Inkscape (v1.3.2).

## Supplemental figures

**Figure S1.**
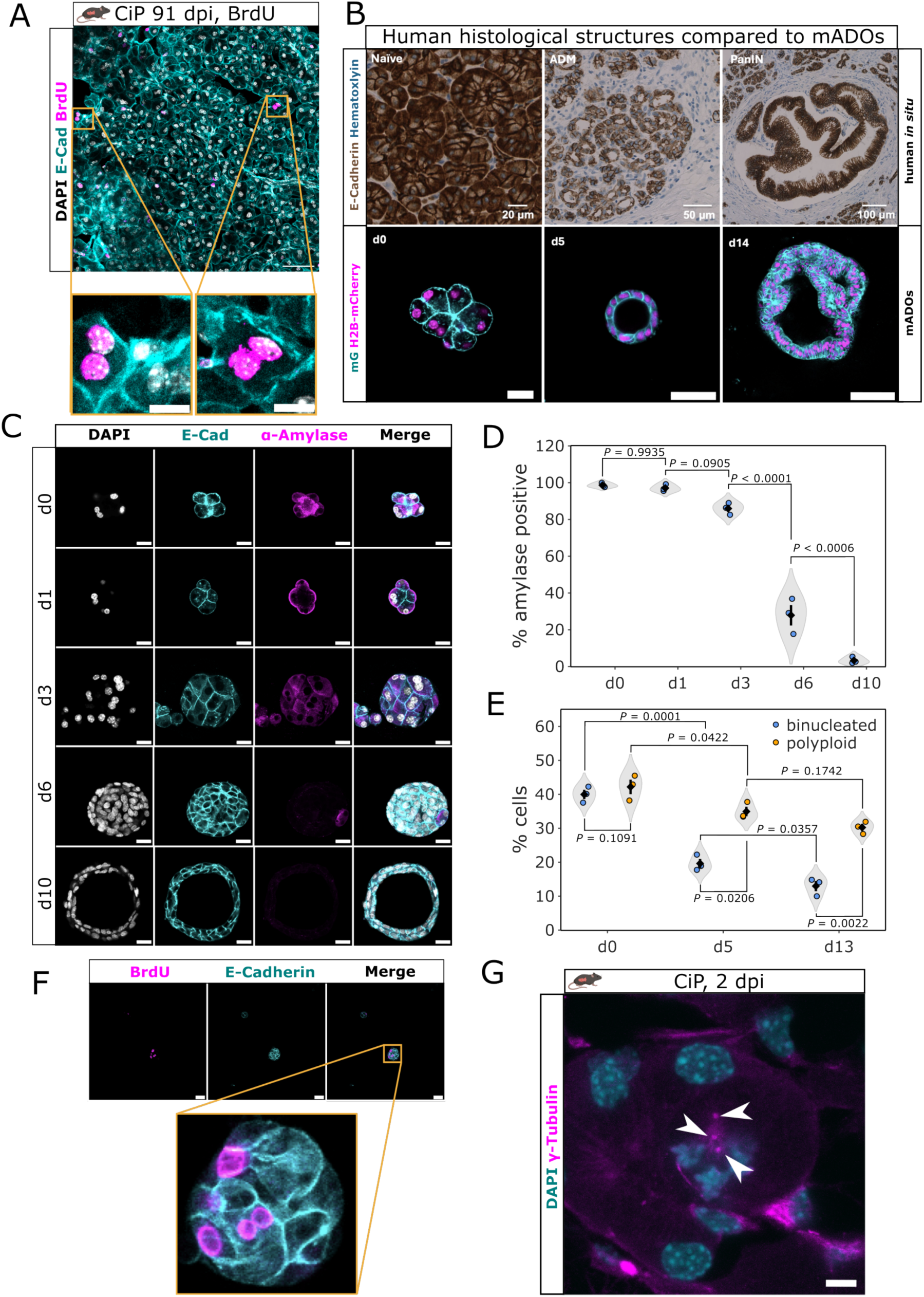
(A) IF image of post-regenerative mouse pancreatic tissue after cerulein-induced pancreatitis (91 dpi) and subsequent BrdU treatment. BrdU pulse was administered at 0 dpi. DAPI (white), E-Cadherin (cyan), BrdU (magenta). Yellow squares indicated zoomed-regions highlighting binucleated cells with BrdU-positive nuclei. Scale bar 50 μm. (B) Top row: immunohistochemical images of human pancreas tissue sections showing naïve, ADM and PanIN-lesion tissue. E-Cadherin (brown), hematoxylin (blue). Scale bar lengths indicated on images. Bottom row: IF images of mouse pancreatic acinar-derived organoids (mADOs). mADO development recapitulates ADM and PanIN formation *in vitro*. Scale bar 20 μm (left), 50 μm (center), 100 μm (right). (C) Time series of IF images representing mADO formation at different times in culture (0d, 1d, 3d, 6d, 10d). DAPI (white), E-Cadherin (cyan), Amylase (magenta). Scale bars: 20 μm. (D) Quantification of α-amylase-expressing cells in mADOs at different times in culture. *P* values were calculated by one-way ANOVA followed by Tukey’s post hoc test (n=3 mice). (E) Quantification of binucleated (dark grey) and polyploid (light grey) cells in mADOs at different times in culture. Ploidy was determined based on the image analysis pipeline presented in Figure S3C. *P* values between binucleated and polyploid cells were calculated using a two-sided paired Student’s *t* test for each timepoint (d0, d5, d13). Changes over time were compared using a Repeated Measures ANVOA followed by Tukey’s post hoc test for binucleated and polyploid cells respectively (n=3 mice). (F) IF images of short-term BrdU pulse-chase experiments in mADOs (24h in culture). BrdU-pulse given at 22 h, chase time 2h. BrdU (magenta), E-Cadherin (cyan). Yellow box indicates zoomed region showing a cell cluster with binucleated BrdU-positive cell. Scale bars: 50 μm. (G) IF image of mouse pancreatic tissue after CiP (2 dpi) showing a mitotic polyploid cell. DAPI (cyan), γ-Tubulin (magenta). White arrows indicating centrosome positions. Scale bar: 5 μm

**Figure S2.**
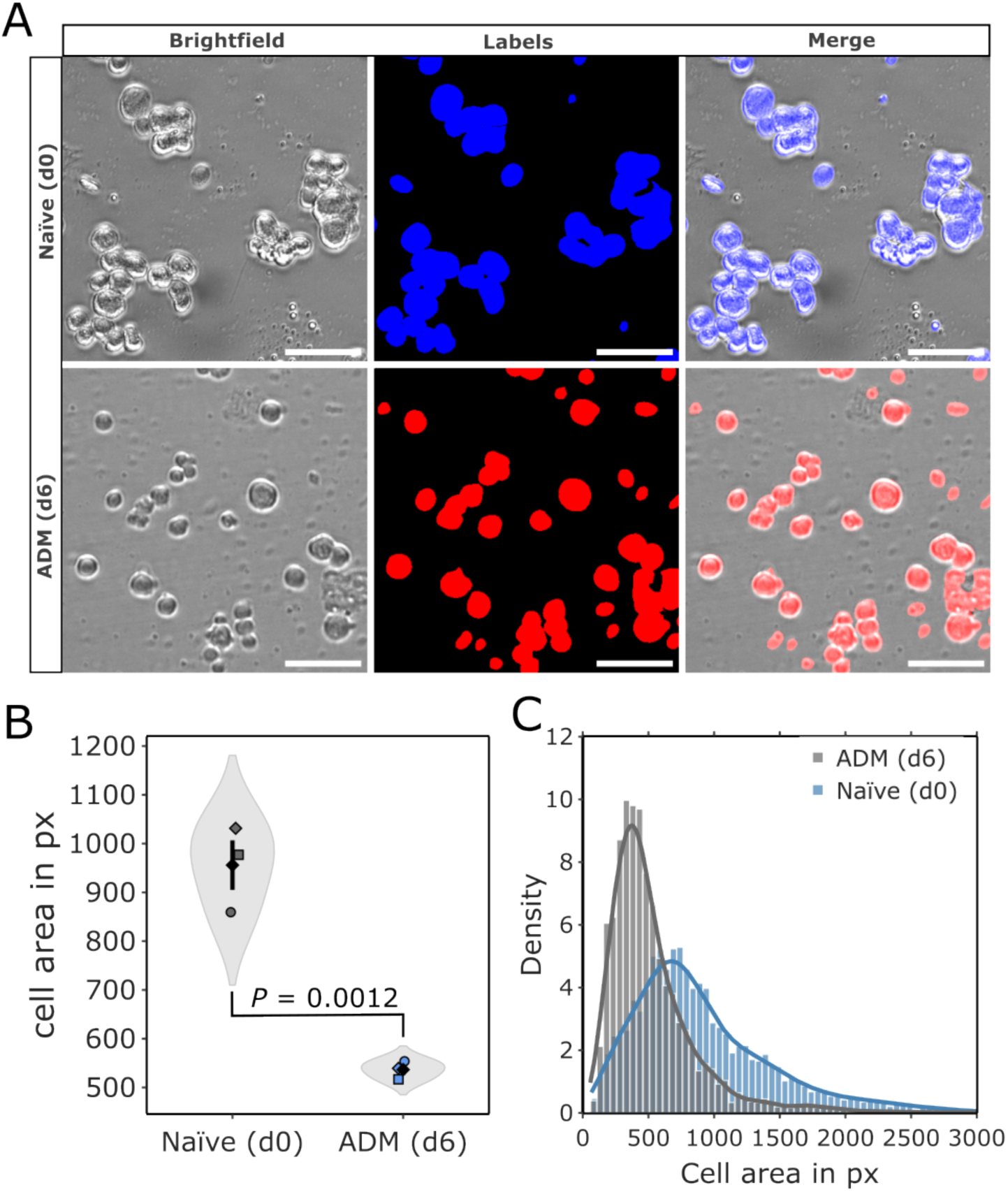
(A) Brightfield images of naïve acinar (top row) and ADM (d6 mADOs, bottom row) cells (left column) with corresponding segmentation masks in blue and red (center column) and overlay (right column) used for cell size measurements. Scale bar 50 μm. (B) Cell area measurements of naïve acinar and ADM cells based on dissociated tissue and mADOs in pixels (px). *P* value was calculated using a two-sided unpaired Student’s *t* test (n = 3 mice). (C) Histogram of cell size distributions from naïve (blue) and ADM (grey) cells.

**Figure S3.**
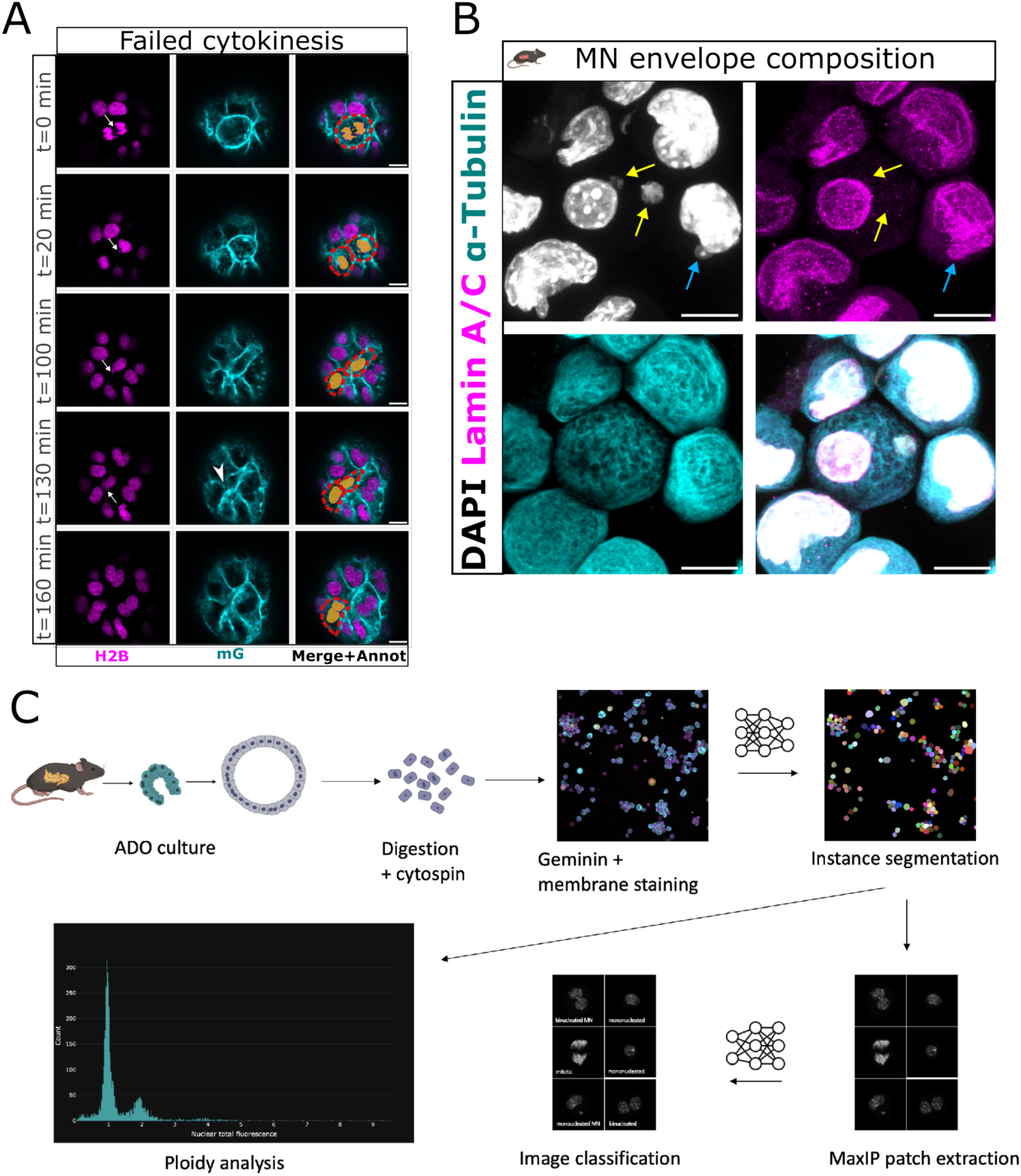
(A) Live-cell image series of a d5 H2B-mCherry/mG mADO cell exhibiting a lagging chromosome with micronucleation (white arrow) and a subsequent failed cytokinesis resulting in hyperpolyploidization. White arrowhead indicates cleavage furrow regression. H2B (magenta), mG (cyan). Yellow annotation indicates nuclei, and red annotation indicates cell borders. Scale bar 10 μm. (B) Immunofluorescence images of dissociated d5 mADOs cells stained to assess micronuclear envelope composition for Lamin A/C. DAPI (white), Lamin A/C (magenta), α-Tubulin (cyan). Scale bar 10 μm. (C) Schematic experimental and image analysis workflow to classify cells based on confocal images and assess ploidy state as well as γH2A.X signal intensities quantitatively. mADO cultures were dissociated into single cells, fixated on microscopy slides using a Cytospin centrifuge and stained for geminin (cell cycle marker) and α-Tubulin (cell body marker) and nuclear staining using DAPI to measure DNA content. Maximum intensity projections (MaxIP) of α-Tubulin confocal images were instance segmented using a custom Cellpose model. Each cell’s MaxIP of the DAPI channel was cropped based on its cell segmentation shape and fed into a ResNet50 convolutional neuronal network to classify it according to either mononucleated, mononucleated/micronucleated, binucleated, binucleated/micronucleated or mitotic. Instance segmentation images were further used to extract various numerical features to correct for mis-segmented cells and assess measures such as DAPI, geminin and γH2A.X signal intensities. Scheme was in parts created in Biorender.

**Figure S4.**
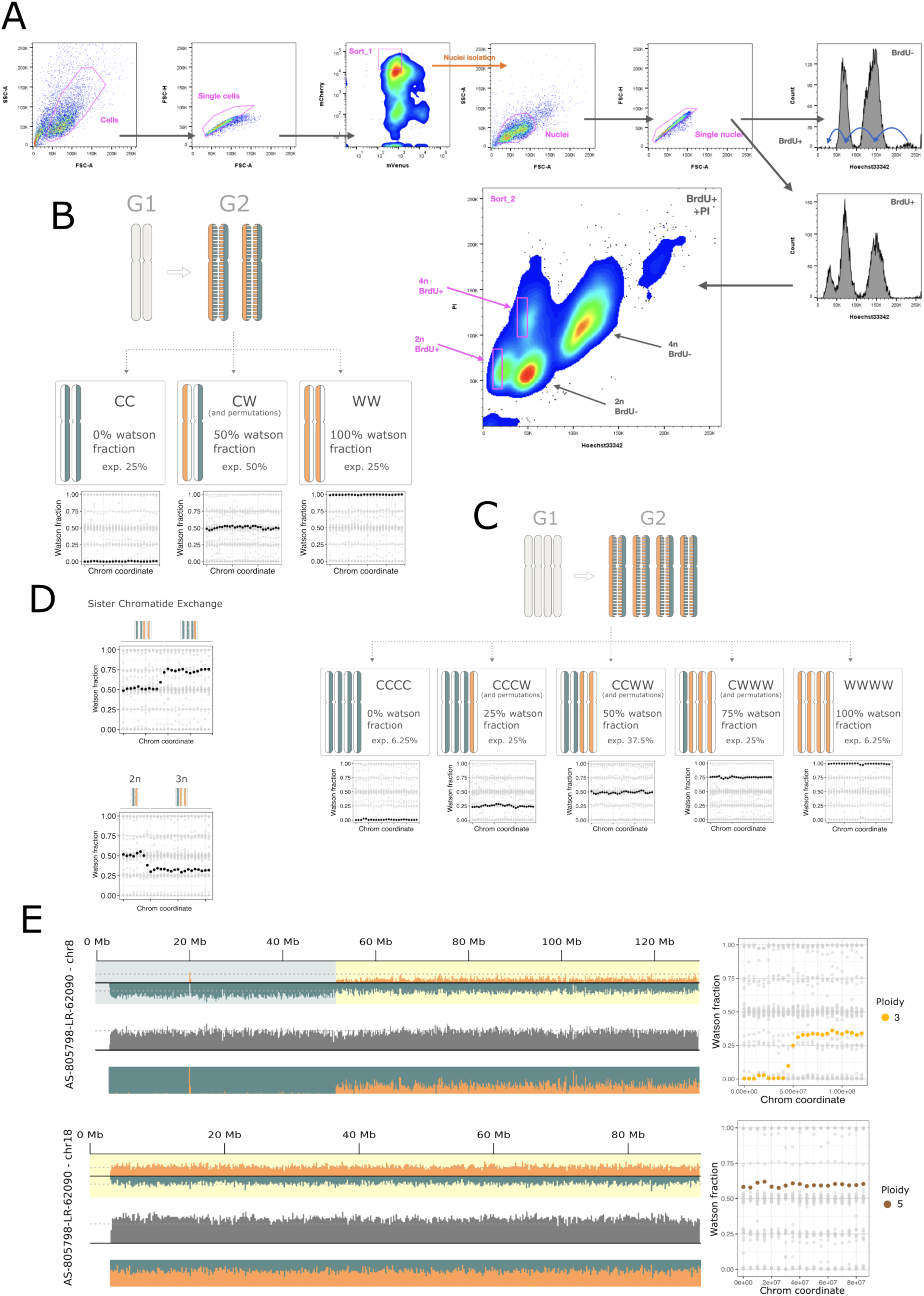
(A) Sorting strategy to sort single nuclei for Strand-seq. An initial sorting step consisted of sorting mCherry^+^ cells from Fucci2, indicating non-cycling cells (sort_1). After lysis, doublet nuclei were excluded and BrdU incorporation was assessed by quenching of Hoechst33342 (Hoechst 33342 signals roughly half upon BrdU incorporation). Blue arrows in the BrdU^−^ (no BrdU during mADO culture) control graph indicate, where the fractions of different ploidy would be expected to shift if BrdU is incorporated. BrdU^+^ graph shows the sorting from a BrdU treated sample. Propidium iodide (PI) is used to separate the Hoechst peaks by ploidy, as PI is not affected by BrdU incorporation and sort BrdU^+^ nuclei for Strand-seq (sort_2). Magenta annotations indicate gating. 2n: diploid nuclei, 4n: tetraploid nuclei, SSC-A: side scatter area signal, FSC-A: forward scatter area signal, FSC-H: forward scatter height signal. (B) Theoretical distributions of Watson fractions (WF) - defined as the proportion of sequencing reads aligned to the Watson strand relative to the total number of aligned reads within a genomic bin - are shown for diploid loci. (C) As (B), but for tetraploid loci. During DNA replication, each strand has a 50% probability to be used as template, resulting in possible configurations of CC, CW, WC, and WW in the daughter cells with diploid chromosomes, with analogous configurations for tetraploid loci. Plotting the WF of genomic bins across chromosomes allows for the inference of cellular ploidy, as some values can only be explained by a specific ploidy/copy number value (within a reasonable range), e.g. 33% and 66% are specific for 3n, 25% and 75% for 4n. Additionally, combining WF data with normalized total read counts enables estimation of chromosome copy number when WFs take ambiguous values such as 0% and 100%. (D) The same type of visualization allows to spot sister chromatid exchanges (top panel) and aneuploidies (bottom panel). (E) Single-cell Strand-seq plot showing a triploid chromosome (chr8) and a pentaploid chromosome (chr18) from a tetraploid cell. The left panels illustrate coverage metrics, including strand-specific read depth, total read depth, and normalized Watson/Crick fractions at a 200 kb bin resolution, with dotted lines indicating the median value for the cell. The right panels show WF distributions along chromosomal coordinates, with data points color-coded by ploidy. Gray background points represent corresponding chromosome data from other cells. WFs are aggregated in 5 Mb intervals and smoothed over a 10 Mb window.

**Figure S5.**
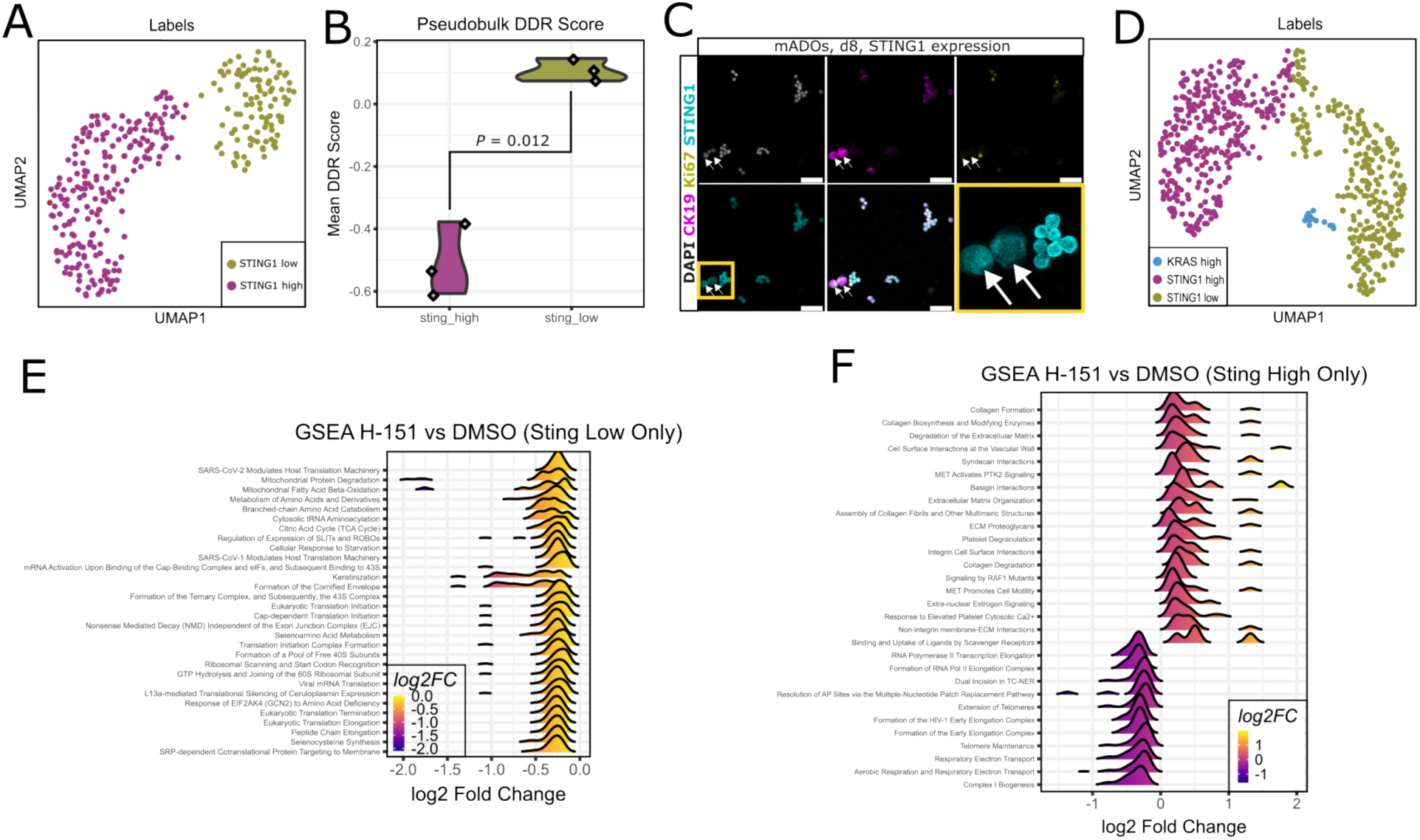
(A) UMAP plots of single-cell proteomic data indicating DMSO-treated cells displaying ploidy and cluster labels (defined by STING1 protein expression, as show in Figure 5A). (B) DNA damage and repair score of pseudobulks between the two clusters. (C) Representative IF image of d8 mADO cells stained for STING1 (cyan), CK19 (magenta), Ki67 (yellow and DAPI (white). Yellow magnification box highlights STING1 low polyploid cells (white arrows) next to STING high diploid cells. Scale bar 50 μm. (D) UMAP plots of DMSO-treated and H-151-treated cells displaying ploidy and cluster labels, defined by STING1 and KRAS expression (as shown in Figure 5G). (E) Ridge plots displaying the kernel density curves of log2 fold changes (log2FC) for genes within each of the top 30 Reactome pathways identified by GSEA. Comparison shows H-151 versus DMSO treatment in the STING1-low cluster (F) As (E), but log2FC are shown for the comparison between H-151 and DMSO for the STING1-high cluster.

**Figure S6.**
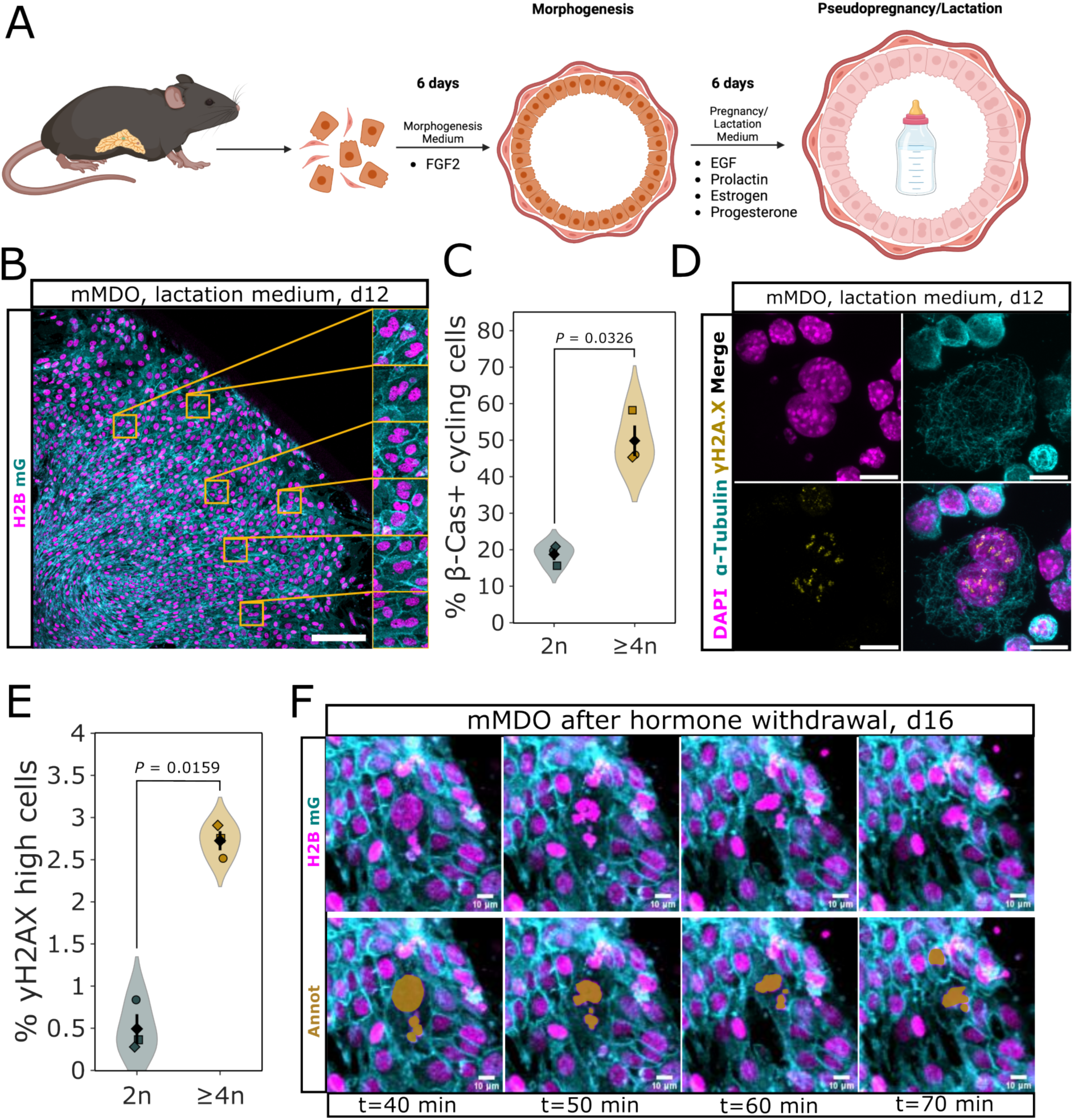
(A) Experimental workflow to generate lactating mouse mammary gland organoids (mMDOs) consisting of diploid and polyploid secretory cells. The procedure includes cells extraction, 6 days morphogenesis supplementing with FGF2 and 6 days of pseudopregnancy/lactation including polyploidization and milk production induced by EGF, prolactin, estrogen and progesterone. Scheme was created in Biorender. (B) Image from live-cell microscopy of H2B-mCherry/mG mMDOs after adding lactation medium. Yellow boxes highlight regions showing binucleated cells. H2B (magenta), mG (cyan). Scale bar 50 μm. (C) Quantification of β-Casein-positive cycling cells among the diploid and polyploid fraction in d12 mMDOs (6 days MOM + 6 days PLM). *P* value was calculated using a two-sided paired Student’s t test (n = 3 mice). (D) Immunofluorescence images of γH2A.X high binucleated mMDO cell with micronuclei. DAPI (white), α-Tubulin (cyan), γH2A.X (yellow). Scale bar 10 μm. (E) γH2A.X measurement of diploid and polyploid mMDO cells at d12 after morphogenesis and pregnancy/lactation hormone treatment. *P* value was calculated using a two-sided paired Student’s *t* test (n = 3 mice). (F) Live-cell image series of an H2B-mCherry/mG mMDO after hormone withdrawal showing a micronucleated cell undergoing mitosis. Upper row: H2B (magenta), mG (cyan). Bottom row: yellow annotation indicated nuclei of interest. Scale bar 10 μm.

**Table S1:** List of used antibodies.

| Antibody | Source | Identifiers | Dilution/Concentration |
| --- | --- | --- | --- |
| **Rat anti-BrdU** | Abcam | Cat# ab6326 | 1:100 |
| **Rat anti-Cytokeratin 19** | DSHB | TROMA-III (conc.) | 3 µg/ml |
| **Rabbit anti-DCLK1** | Abgent | Cat# AP7219b | 1:200 |
| **Rat anti-E-Cadherin** | Thermo Fisher | Cat# 13-1900 | 1:1000 |
| **Rabbit anti-Geminin** | Abcam | Cat# ab195047 | 1:200 |
| **Mouse anti-Lamin A/C** | Cell Signaling | Cat# 4777S | 1:200 |
| **Rabbit anti-Pericentrin** | Abcam | Cat# ab4448 | 1:100 |
| **Rabbit anti-phospho histone H3** | Merck | Cat# 06-570 | 1:250 |
| **Rabbit anti-STING1** | Invitrogen | Cat# PA5-23381 | 1:100 |
| **Rat anti-Tubulin** | Abcam | Cat# ab6160 | 1:1000 |
| **Rabbit anti-α-Amylase** | Sigma Aldrich | Cat# AB273 | 1:100 |
| **Mouse anti-β-Casein** | Santa Cruz Biotechnology | Cat# sc-166530 | 1:200 |
| **Mouse anti-γ-Tubulin** | Abcam | Cat# ab11316 | 1:100 |
| **Mouse anti-γH2A.X** | Sigma Aldrich | Cat# 05-636 | 1:200 |

### Captions for Movies S1 to S9

**Movie S1.** Maximum intensity projection time lapse movie of H2B-mCherry/mG mADO at d4 highlighting binucleated cell division. Left: H2B-mCherry channel (magenta) with bounding box (yellow) and nuclei annotation (blue) of mitotic binucleated cell. Right: overlay of H2BmCherry (magenta) and mG (cyan) channels with yellow bounding box indicating mitotic binucleated cell. Scale bar 30 μm.

**Movie S2.** Maximum intensity projection time lapse movie of H2B-mCherry mADO at d8 highlighting binucleated cell multipolar division. H2B-mCherry channel: magenta, nuclei annotation of mitotic binucleated cell: blue. Scale bar 10 μm.

**Movie S3.** Maximum intensity projection time lapse movie of EGFP-Tuba (cyan) mADO at 12 showing binucleated cell with nuclei oriented along the short axis and cell division along the long axis. Scale bar 10 μm.

**Movie S4.** Maximum intensity projection time lapse movie of EGFP-Tuba (cyan) mADO at 12 showing binucleated cell with nuclei oriented along the long/division axis. After initial multipolar spindle formation, the cell divides in a bipolar manner. Scale bar 10 μm.

**Movie S5.** Maximum intensity projection time lapse movie of EGFP-Tuba (cyan) mADO at 12 showing binucleated cell with nuclei oriented along the long cell axis dividing in a multipolar manner. Scale bar 10 μm.

**Movie S6.** Maximum intensity projection time lapse movie of H2B-mCherry (magenta) mADO directly after cell isolation (d0) highlighting binucleated cell divisions with yellow bounding boxes. Scale bar 10 μm.

**Movie S7.** Maximum intensity projection time lapse movie of H2B-mCherry (grey) mADO at d13 showing a proliferative micronucleated cell with repeated mitotic errors. Scale bar 20 μm.

**Movie S8.** Maximum intensity projection time lapse movie of H2B-mCherry/mG mMDO at d16 showing binucleated cell multipolar division. H2B-mCherry channel: magenta, mG channel: cyan. Scale bar 10 μm.

## Author Contributions

Conceptualization: A.M.V. and J.B. Writing, review and editing: J.B., A.C., E.P. and A.M.V. Methodology: J.B., Sa.A., So.A., A.S. and D.K. performed the in vitro and in vivo experiments regarding pancreatic acinar cells. J.B., V.E.L. and S.B.C. performed the mammary-derived organoids experiments. J.B performed the live-cell imaging experiments and U.E. advised the live-cell imaging. K.B. and J.P.M. did the library preparation for the Strand-seq experiment. J.B. performed the Strand seq sequencing procedure. A.S.A. and J.K. designed and conducted the single-cell proteomics experiment. Formal imaging data analysis: J.B. Formal sequencing and single-cell proteomics data analysis: E.F.; Si.A. advised the data analysis of the single-cell proteomics data. Funding acquisition: A.M.V.

## Acknowledgements

We would like to thank the lab of Jan Korbel, especially Maja Starostecka and Marco Raffaele Cosenza for advice on the Strand seq protocol and technical support on the cell sorting and data analysis. We additionally thank Jan Ellenberg for providing the H2B-mCherry/mG mouse line, Takashi Hiragi for providing the EGFP-Tuba mouse line and Michael Milsom for providing the Fucci2 mouse line. Furthermore, we thank Axel Roers for providing the H-151 STING inhibitor. This work was supported by the DKFZ-MOST Cooperation Program, the SFB873 of the German Research Foundation, and the DKFZ Heidelberg.

