## Supplementary material for "Inflammatory injury drives CIN and STING loss in polyploid pancreatic acinar cells": Data-S1

### Data-S1 – Results

##### Quality Control and Characterization of Single-Cell Proteomics Data

We performed single-cell proteomics (SCP) on 734 pancreatic derived organoid cells equally distributed between wild-type (DMSO, n=367) and STING inhibitor-treated (H-151, n=367) conditions, with cells sorted into three ploidy groups (2n, 4n, 8n) across two 384 well plates (Data-S1 Figure 1A). Our SCP pipeline yielded high-quality data with a median of approximately 4,000 protein identifications per cell (Data-S1Figure 1B) and 6,100 total unique proteins across the entire dataset. Protein detection showed 98% overlap between conditions (Data-S1Figure 1C), with only 37 DMSO-specific and 77 H-151-specific proteins, indicating that STING inhibition does not majorly deform the proteome. Protein intensity distributions were nearly identical between conditions (Data-S1Figure 1D), confirming comparable detection sensitivity and data quality across experimental groups. To evaluate the technical reproducibility and stability of mass spectrometry for SCP, we monitored total protein intensity across cells in their chronological processing order which showed total protein intensity remained stable across all runs (Data-S1Figure 1E), with no ion suppression, or carryover effects. Total intensity per cell was consistent across ploidy states (Data-S1Figure 1F), demonstrating that the workflow samples cellular proteomes uniformly regardless of cell size. Plate-to-plate comparison revealed highly similar intensity distributions (median log10 ~5.7-5.8) (Data-S1Figure 1G), validating reproducible sample preparation and instrument performance. Quantitative assessment of relative standard deviation (RSD) showed that within-plate variability (median ~75%) was substantially higher than between-plate variability (Data-S1Figure 1H), confirming that biological heterogeneity dominates over technical batch effects. Principal component analysis demonstrated that cells from different plates were intermixed in PC space (Data-S1Figure 1I), with PC1 capturing 29.4% and PC2 capturing 5.8% of variance. The absence of plate-specific clustering confirms absence of batch effects, while the variance distribution across the first five PCs (Data-S1Figure 1J) follows the expected pattern for biological datasets with dominant factors (29%, 6%, 4%, 3%, 2%). Together, these quality metrics demonstrate that our single-cell proteomics workflow achieves high technical reproducibility, comprehensive proteome coverage, and robust quantitative performance suitable for detecting biologically meaningful differences between experimental conditions.

### Data-S1 — Figures

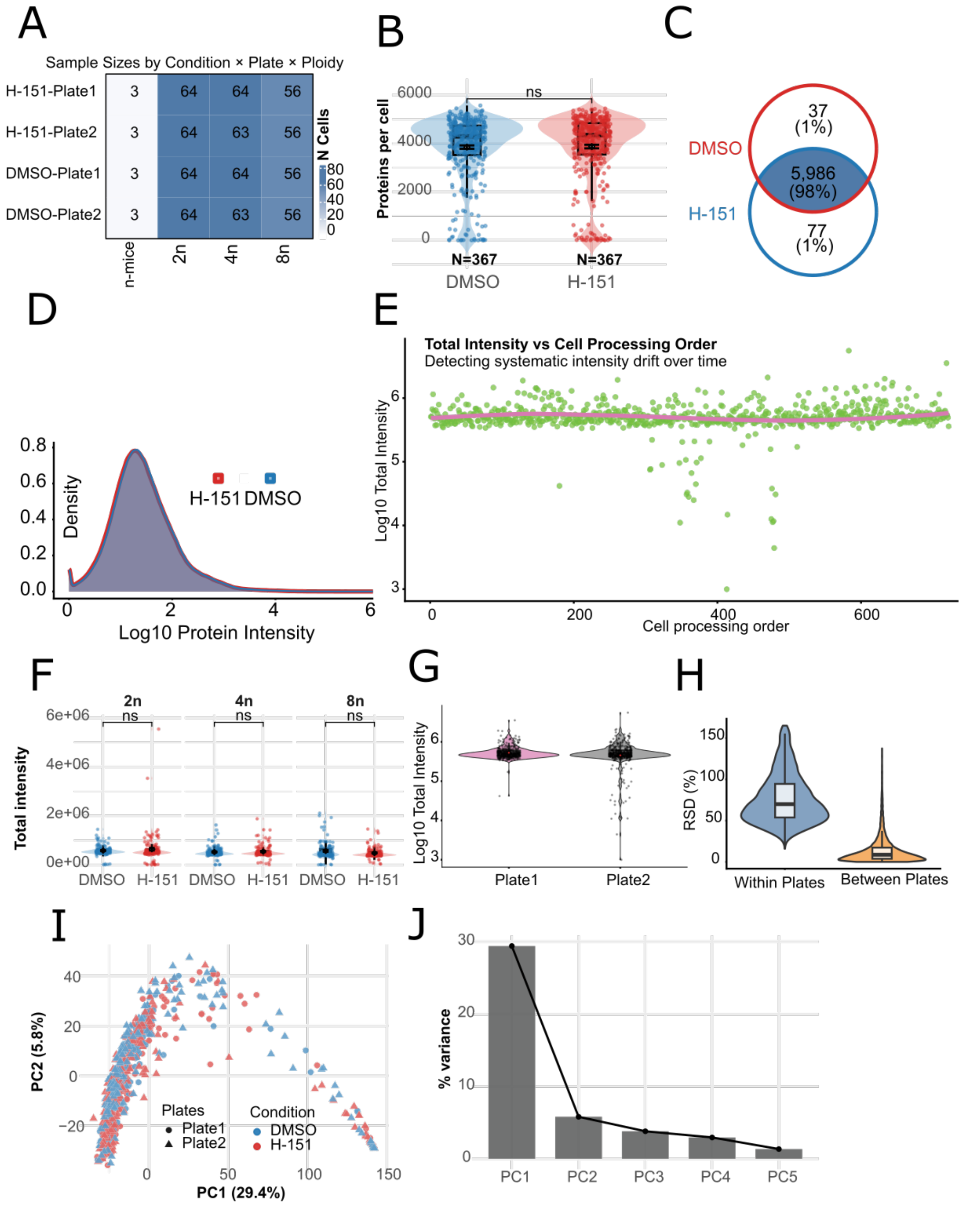


**Data-S1 Figure 1. Quality control and sample characterization of single-cell proteomics data.**

(A) Heatmap showing the distribution of 734 single cells across experimental groups: two conditions (DMSO, H-151), two plates (Plate1, Plate2), and 3 mice with three ploidy states (2n, 4n, 8n). Numbers indicate cell counts per group.

(B) Distribution of protein identifications per cell for DMSO (N=367) and H-151 (N=367) conditions. Violin plots show kernel density estimates with overlaid boxplots (center line: median, box: interquartile range) and individual cell measurements (dots). ns: not significant by Wilcoxon test.

(C) Venn diagram showing shared (5,986 proteins, 98%) and condition-specific proteins between DMSO and H-151. Percentages calculated from total detected proteins (6,100).
(D) Density distribution of protein intensities across all proteins and cells for DMSO (blue) and H-151 (red) conditions.
(E) Total protein intensity plotted against cell processing order with locally estimated scatterplot smoothing (LOESS) smoothing lines.
(F) Total protein intensity per cell by ploidy (2n, 4n, 8n) and condition (DMSO, H-151). Violin plots show kernel density estimates with overlaid boxplots and individual cell measurements (dots). ns: not significant by Wilcoxon test.
(G) Violin plots comparing total protein intensity distributions between technical replicate plates.
(H) Violin plots comparing relative standard deviation (RSD) of protein expression within plates versus between plates.
(I) Principal component analysis (PCA) of single-cell proteomes showing the first two principal components (PC1: 29.4% variance, PC2: 5.8% variance). Cells are colored by condition (blue: DMSO, red:H-151) and shaped by plate (circles: Plate1, triangles: Plate2).
(J) Scree plot showing variance explained by the first five principal components.
